# A human cell-free translation screen identifies the NT-2 mycotoxin as a ribosomal inhibitor that binds the peptidyl transferase center

**DOI:** 10.1101/2025.10.11.680285

**Authors:** Nino Schwaller, Dominic Andenmatten, Jonas Luginbühl, Julius Rabl, Helena Baur, Marc Chambon, Jonathan Vesin, Gerardo Turcatti, Evangelos D. Karousis

**Affiliations:** Department of Chemistry, Biochemistry and Pharmaceutical Sciences, University of Bern, Bern, 3012, Switzerland; Multidisciplinary Center for Infectious Diseases, University of Bern, Bern, 3012, Switzerland; ETH Zurich, Cryo-EM Knowledge Hub, 8093 Zurich, Switzerland; Institute of Cell Biology, University of Bern, Bern, 3012, Switzerland; Biomolecular Screening Facility, EPFL, Lausanne, Switzerland

**Keywords:** Cell-free translation, high-throughput screening, protein synthesis inhibitors, mycotoxin, mRNA translation, NT-2 toxin, ribosome, translation regulation, Fusarium, peptidyl transferase center, SERBP1, dormant ribosomes

## Abstract

Translation inhibitors are invaluable for probing ribosome function and therapeutic applications, but systematic discovery in human systems is limited by the lack of scalable, screening-compatible cell-free platforms. Here, we establish a robust high-throughput screening using human lysates that bypasses cellular cytotoxic effects. After screening ∼28,000 small molecules, we identified known and a novel translation inhibitor, including NT-2, a trichothecene mycotoxin produced by the pathogenic *Fusarium sporotrichioides*. NT-2 suppressed protein synthesis in human cells and yeast lysates, while sparing translation in bacteria and intact yeast cells. Cryo-EM at 1.76 Å revealed NT-2 bound at the peptidyl transferase center of the human 60S ribosome, confirming NT-2 as a ribosomal inhibitor associated with ribosomes in an inactive eEF2/SERBP1-associated dormant state. Together, these results expose NT-2 as a previously unrecognized environmental inhibitor of mammalian protein synthesis and demonstrate the power of cell-free translation screening to reveal new inhibitors with unexpected ribosome fates.

## Introduction

Despite the central role of protein synthesis and translational control in disease and therapy, the discovery of new translation inhibitors is challenging. Screenings using living cells have uncovered translation-modulating compounds ^1–3^ but cell-based screenings are inherently limited by compound uptake, cytotoxicity, and physiological complexity, which can obscure target-specific effects ^4,5^. False positives and overlooked candidates remain a persistent problem, highlighting the need for complementary platforms with high mechanistic resolution ^6^ and they often fail to discriminate between cell growth inhibitors from those acting on specific mRNA translation mechanisms, such as IRES- or RAN-mediated translation ^7,8^. While useful, traditional cell-free systems such as rabbit reticulocyte lysate or *E.coli* lysates lack human-specific context, they present tissue-specific artefacts and are labour-intensive to prepare at scale ^9–11^. Moreover, they are generally incompatible with high-throughput screening formats, some enzymatic assays, and do not support physiologically relevant post-translational processes ^6,9,12^.

This gap is particularly evident for natural toxins, where food-borne contaminants can pose serious health hazards. Mycotoxins are pervasive fungal metabolites that contaminate global food production and exert potent bioactivities, often targeting core cellular processes ^13,14^. Trichothecenes are toxic fungal metabolites produced by species of the genus *Fusarium* that increasingly contaminate grains due to changing climate and agricultural practices ^15^. They can cause a broad spectrum of health effects, including gastrointestinal, immunological, neurological, dermal, and reproductive toxicity, with some showing carcinogenic potential in animal models ^14,16^. Globally, around one-quarter of crops exceed regulatory thresholds for mycotoxin contamination, and 60–80% contain detectable levels ^16^. These risks explain the urgent need to characterize, monitor, and mitigate trichothecene contamination in food and feed supplies. Several Trichothecenes can cause eukaryotic translation inhibition ^17–20^. Although traditionally studied in the context of food safety, their potent effects make them promising candidates for therapeutic development, especially in antiviral ^21^ and anticancer contexts ^22,23^. Yet few fungal metabolites beyond classical trichothecenes have been systematically examined in human systems ^16^. Structural studies have shown that trichothecene toxins bind the ribosomal peptidyl transferase center and inhibit translation^24^.

NT-2 (15-deacetyl-neosolaniol) is a Type A trichothecene mycotoxin first isolated from *Fusarium sporotrichioides* ^15,25,26^. NT-2 carries an epoxide moiety at C12–13, a feature essential for trichothecene bioactivity, but its unique substitution pattern, including multiple hydroxyl groups, chemically distinguishes it from the previously studied T-2 and HT-2 toxins^14,24^.

Here, we establish the first high-throughput cell-free translation screening (CFTS) using human lysates, fully compatible with 384-well formats. Screening ~28,000 compounds revealed NT-2, a previously uncharacterized trichothecene mycotoxin from *Fusarium*, as a potent inhibitor of eukaryotic translation. Cryo-EM at 1.76 Å placed NT-2 in the peptidyl transferase center (PTC) and revealed an accumulation of ribosomes in an inactive eEF2/SERBP1-associated dormant state. Together, these findings reveal an overlooked environmental threat to human translation and demonstrate the power of CFTS to uncover both new inhibitors and unexpected ribosome fates.

## Results

### Development of a human Cell-Free Translation Screening Platform

To overcome the limitations of living cells in high-throughput discovery, we established a massively parallel cell-free translation screening based on human dual-centrifugation lysates ^12,27,28^. Cytoplasmically enriched extracts preserve key human translation features and provide sufficient material for thousands of reactions ^27^. As illustrated in Fig. 1A, the workflow involves lysate preparation, the addition of reporter mRNAs and compounds in 384-well plates, and automated luminescence readout, enabling compatibility with large-scale screening.

**Figure 1.**
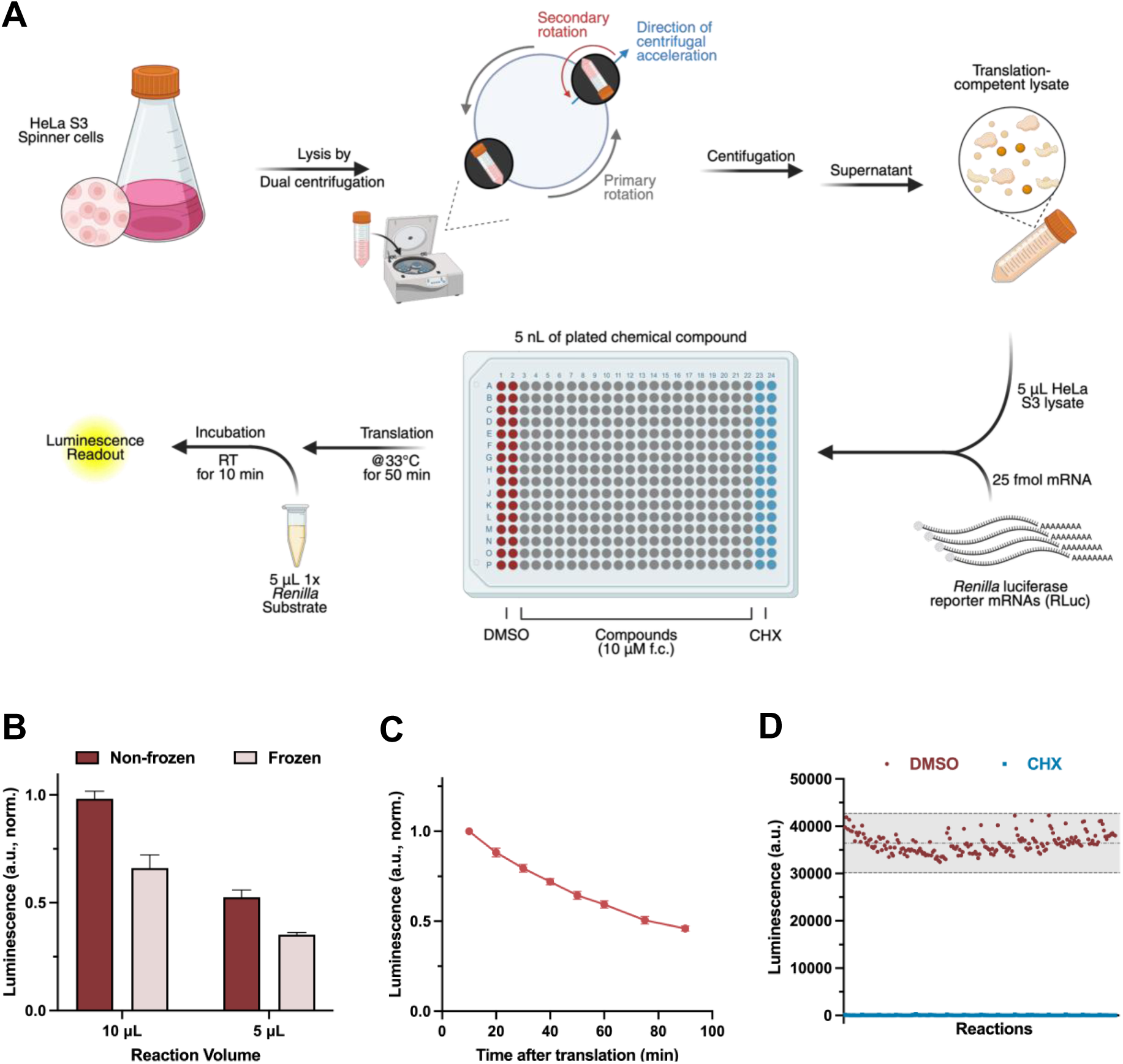
CFTS platform development and optimization. **(A)** CFTS workflow. Dual centrifugation-produced HeLa S3 lysates are combined with *Renilla* luciferase reporter mRNAs and dispensed into 384-well plates preloaded with compounds. Translation was quantified by luminescence. **(B)** Luminescence output from reactions performed at 5 μL and 10 μL volumes using freshly assembled or pre-frozen reaction mixes. **(C)** Luminescence signal stability over 90 minutes following substrate addition. **(D)** Z’ factor assessment with DMSO (vehicle) or cycloheximide (CHX, positive control) with mean ± 3SD shown as dotted lines. For B and C, data are presented as mean values of three biological replicates (sets of translation reactions) averaged after three measurements ± SD.

Translation of capped and polyadenylated Renilla luciferase mRNA provides a quantitative readout for protein synthesis, and pre-spotted compounds reduce well-to-well variation. To maximize throughput without compromising fidelity, we optimized several parameters. Low reaction volumes (5 μL per well) retain signal intensity, lysate-mRNA mixes can be frozen and thawed with a minor reduction in activity (Fig. 1B), and luminescent signals remain stable within the time window of 20 minutes required to process more than 10 plates in parallel. (Fig. 1C). Assay quality was benchmarked with cycloheximide, yielding a Z′ factor of 0.82, reflecting a dynamic range exceeding three orders of magnitude and a clear separation between uninhibited and fully inhibited reactions ^29^ (Fig. 1D). The assay displayed robust performance across screening plates with minimal inter-plate variability (coefficient of variation < 5%; Sup. Table 1). These performance metrics establish CFTS as a robust and human-specific platform for scalable discovery of translation inhibitors.

### Identification of active compounds in chemically diverse libraries

We performed the high-throughput CFTS to evaluate 27,763 small molecules from six mechanistically and chemically diverse libraries for their ability to modulate human translation. Compounds were tested at a final concentration of 10 µM, and those with a hit score above 0.6 were classified as primary active hits. The hit score was calculated from the luminescence signal relative to uninhibited (DMSO) and fully inhibited (CHX) controls, where a value of 1 corresponds to complete inhibition equivalent to CHX and 0 corresponds to no inhibition (DMSO).

The Prestwick Library (Fig. 2A), made up of 1280 FDA/EMA/PMDA-approved drugs with noted bioactivity and the kinases inhibitor collection (275 compounds), yielded 17 active compounds out of the 1,555 molecules, including several known translation inhibitors. In the Repurposing Library (Fig. 2B), which contains molecules that have reached clinical or preclinical development, 25 actives were identified from 4,477 compounds, with puromycin among the top hits, serving as a positive internal control. The Protein–Protein Interaction (PPI) Library (Fig. 2C), which contains compounds that interfere with known protein-protein interactions, showed the highest apparent hit rate, with 277 actives among 2,156 compounds screened. Seven scored above 0.9, likely indicating enrichment of molecules targeting interaction surfaces essential for ribosome assembly or factor recruitment. To select the most active compounds, we applied a stringent cut-off of 0.7, reducing the primary hits to 90.

**Figure 2.**
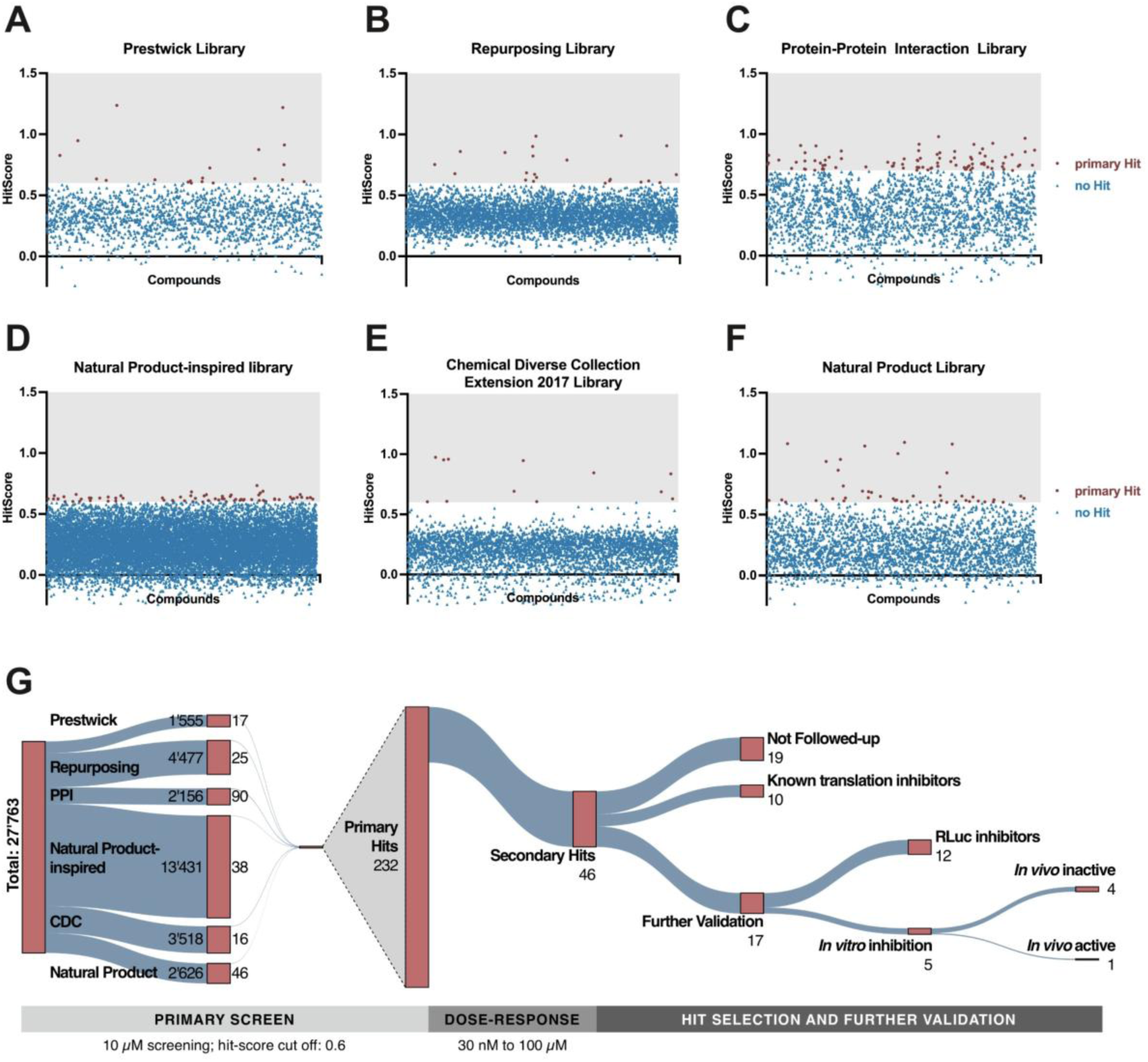
High-throughput identification of translation modulators across chemically diverse libraries. **(A-F)** Compounds with hit scores > 0.6 are positive hits marked with red dots, and negative hits with a score < 0.6 with blue triangles. Hit score distributions from six compound libraries screened by CFTS. Prestwick and Repurposing libraries yielded ∼1% actives, while the PPI library showed the highest density of actives. The Natural product collection contributed multiple top-scoring hits. Compounds were tested at 10 µM in the primary screen (two measurements per compound). Selected hits were validated in independent duplicate dose-response assays. **(G)** Sankey diagram illustrating the sequential filtering of compounds in a multi-step screening workflow. Node widths are proportional to the number of compounds progressing at each stage.

The Natural Product-Inspired Library (Fig. 2D), comprising 13,431 synthetic analogs of natural product scaffolds, yielded 38 actives, though none exceeded a score of 0.75. The Chemically Diverse Collection Extension (Fig. 2E), representing a broad synthetic scaffold space, yielded 16 hits from 3,518 molecules, including four with scores approaching 1.0. Finally, the Natural Products Library (Fig. 2F), comprising 2,626 plant and microbial extracts, yielded 46 primary hits, several with scores greater than 1.0, illustrating the richness of natural metabolites as translation inhibitors. Overall hit rates varied across libraries, ranging from ∼1% in Prestwick and Repurposing to over 10% in the PPI library, highlighting the CFTS capacity to resolve mechanistically diverse modulators across distinct compound classes.

To validate primary hits and assess potency, we performed dose-response assays (30 nM to 100 µM) on 232 top-scoring candidates. Compounds showing reproducible, saturable inhibition curves were classified as secondary hits, yielding 46 molecules, of which 36 had not previously been linked to translation inhibition (Sup. Table 2). To distinguish genuine translation inhibitors from luciferase-specific artefacts, we measured Rluc protein levels by Western blot. Twelve compounds were classified as false positive luciferase inhibitors because the luminescence decrease was not matched by reduced protein levels and persisted when added after translation had completed (Sup. Fig. 1, Sup. Table 2). Ten established inhibitors, including cycloheximide, puromycin, and T-2 toxin, served as internal benchmarks and confirmed assay fidelity (Sup. Table 3). In addition, five molecules were validated as bona fide translation inhibitors (Sup. Table 2). A Sankey diagram summarizes compound filtering through the entire workflow (Fig. 2G).

Together, these results validate CFTS as a high-resolution, scalable platform that identifies modulators of human translation and reveals broader activity patterns across compound classes.

### Validation of the Mycotoxin NT-2 as a translation inhibitor

Among the five compounds confirmed as translation inhibitors after secondary validation, NT-2 was selected for detailed characterization because it belongs to the trichothecene family of *Fusarium* mycotoxins, a group of environmental contaminants with well-documented impacts on food safety and human health. Notably, several trichothecene mycotoxins previously reported to inhibit translation like the T2-toxin and Anguidine ^24,30^ were also identified among the screening hits, further supporting the sensitivity and specificity of the platform (Sup. Table. 5). Our initial human cell-free translation screen identified the mycotoxin NT-2 (15-Deacetylneosolaniol, CAS 76348-84-0) as a translation modulator with a hit score of 0.73. Some trichothecenes are well-established ribosome inhibitors ^26^, yet NT-2 itself remains uncharacterized in this regard. Structurally, NT-2 features a unique combination of substitutions within the trichothecene scaffold, including a C-8 hydroxyl group and a 4β-acetoxy moiety (Fig. 3A), which distinguishes it from better-characterized toxins such as T-2, HT-2, or deoxynivalenol. Secondary validation by dose-response analysis revealed a sigmoidal inhibition curve, with near-complete suppression of protein synthesis at concentrations greater than 30 µM and partial inhibition at lower doses (Fig. 3B). To exclude direct interference with luciferase enzyme activity, we tested NT-2 in a translation assay using a FLAG-tagged RLuc reporter mRNA. Luminescence decreased in a dose-dependent manner, yielding an IC_50_ of 4.1 µM, and this pattern was confirmed by Western blot analysis of FLAG–RLuc protein (Fig. 3C–D). Together, these assays validate NT-2 as a bona fide inhibitor of human protein synthesis.

**Figure 3.**
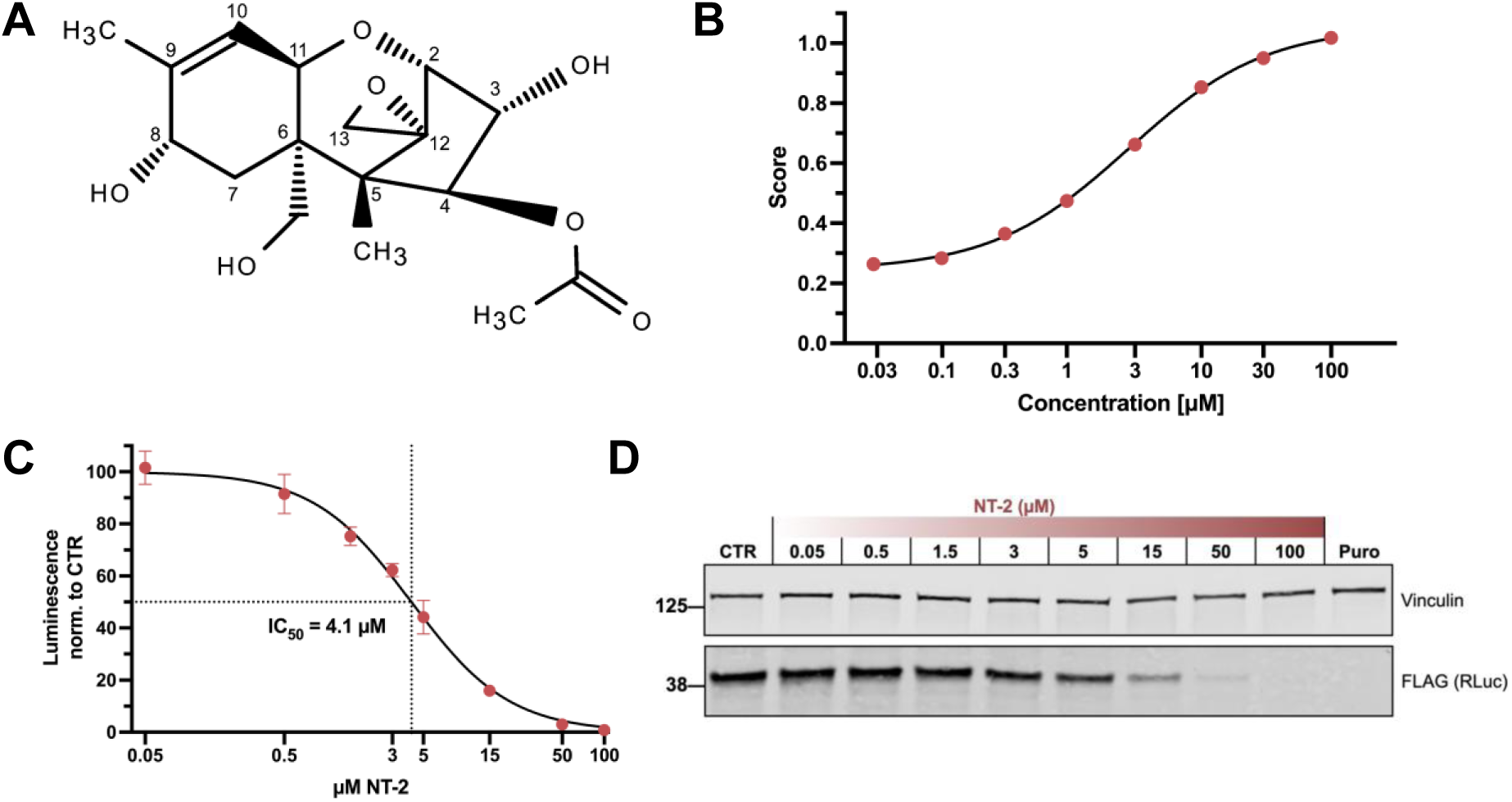
Validation of NT-2 as a translation inhibitor in human cell-free lysates. **(A)** Chemical structure of NT-2, represented in Lewis format **(B)** Dose-response curve from secondary assays showing NT-2–mediated inhibition of *Renilla* luciferase translation. Concentration between 30 nM and 100 µM NT-2 were tested and the inhibition score was calculated, where a value of 0 is no inhibition and 1 is a full inhibition equivalent to CHX. Data are means of two biological replicates (dots) ± SD. **(C)** Cell-free translation reactions were performed using FLAG-tagged *Renilla* luciferase mRNA in HeLa S3 lysate with increasing NT-2 concentrations (50 nM to 100 μM). The measured luminescence was normalized to the negative control containing no supplement (CTR). The calculated IC_50_ for NT-2 was 4.1 μM, based on the average of three biological replicates (dots) ± SD. **(D)** Western blot confirmation of NT-2-mediated translation inhibition. Samples from the reactions in (C) were analyzed by SDS-PAGE and immunoblotting for FLAG-tagged *Renilla* luciferase (translation product) and Vinculin (loading control).

### Eukaryote-specific translation inhibition by NT-2

Because the screening and initial dose-response validation were performed in HeLa S3 lysates, we next asked whether NT-2 also inhibits translation in intact mammalian cells. For cellular assays we used HEK293T cells, to address whether the inhibitory activity identified in HeLa S3 lysates is consistent in a distinct human cellular context. Puromycin incorporation assays in HEK293T cells showed potent suppression of translation, with detectable inhibition at 62.5 nM and complete inhibition at 2 µM (Fig. 4A). Quantification of the puromycin signal yielded a dose-response curve with an IC_50_ of 135 nM (Fig. 4B), indicating substantially greater potency in cells than in lysates. This discrepancy suggests that intracellular factors, such as compound uptake or accumulation, may enhance NT-2 activity *in vivo*. Consistent with this observation, puromycin incorporation assays performed in BHK-21 hamster cells similarly revealed strong suppression of translation upon NT-2 treatment (Sup. Fig. 2), indicating that NT-2 inhibits protein synthesis across mammalian systems.

**Figure 4.**
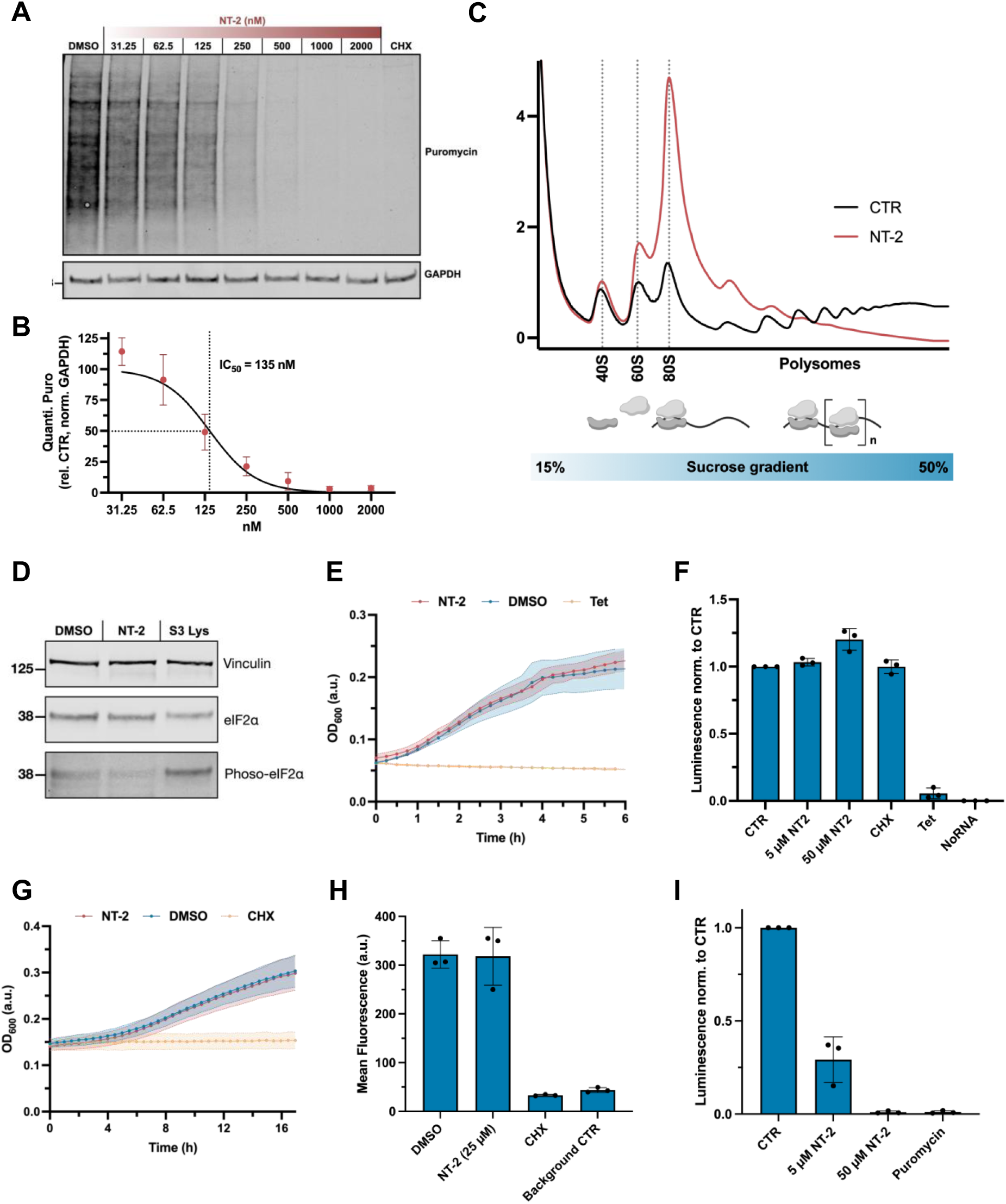
NT-2 activity across human, yeast, and bacterial systems. **(A)** Puromycin incorporation assay in HEK293T cells treated with increasing NT-2 concentrations, DMSO, or CHX. Equal amounts of cell lysate were loaded into a 4-12% Bis-Tris gel and analysed by Western blot probing with anti-puromycin and anti-GAPDH antibodies. **(B)** Quantification of protein synthesis based on signal intensity in the anti-Puromycin-stained blot (Fig 4A) normalized with GAPDH. The calculated IC_50_ for NT-2 in living cells was 135 nM, based on the average of three biological replicates (dots) ± SD. **(C)** Polysome profiles of NT-2-treated HEK293T cells (2 hours with 2 μM NT-2 or DMEM+/+ medium). **(D)** Western blot analysis of eIF2α and phospho-eIF2α in NT-2–treated cells. HEK293T cells were treated with NT-2 (2 μM) or 0.02% DMSO (CTR) for 2 h. Equal amounts of cell lysate were analysed by Western blot probing with anti-Vinculin, anti-eIF2α and anti-phospho-eIF2α antibodies. An equal amount of HeLa S3 lysate was loaded as a positive control for phosphorylation, as described in ^27^. **(E)** Growth curve of *E.coli* treated with 8 µM NT-2 toxin, 0.08% DMSO, or tetracycline (translation-inhibition control). **(F)** Cell-free translation reactions with *Renilla* luciferase mRNA in *E.coli* S30 lysate in the presence of no supplement (CTR), 5 µM NT-2, 50 µM NT-2, cycloheximide (CHX, 100 µg/ml), tetracycline (Tet, 10 µg/ml) or without RNA. The measured luminescence was normalized to the control reaction (CTR). Data are presented as mean ± SD values of three biological replicates (sets of translation reactions) averaged after three measurements. **(G)** Growth curves of *S. cerevisiae* treated with 25 µM NT-2, 0.25% DMSO, or 10 µg/ml CHX as a positive control for translation inhibition. **(H)** Yeast cells were incubated with L-AHA to label newly synthesized proteins and treated with 25 µM NT-2 toxin, 0.25% DMSO (CTR), or 50 µg/ml CHX. Mean fluorescence after click chemistry was quantified by flow cytometry; unlabeled cells indicate background signal. **(I)** Cell-free translation reactions with *Renilla* luciferase mRNA in *S. cerevisiae* lysate in the presence of 5 or 50 µM NT-2 or 1 mg/ml puromycin. The measured luminescence was normalized to the control reaction (CTR) which contained no supplement. Data are presented as mean ± SD values of three biological replicates (sets of translation reactions) averaged after three measurements.

We generated polysome profiles from HEK293T cells treated with 2 µM NT-2 to further investigate the mechanism of translation inhibition. In DMSO-treated cells, prominent polysome and balanced 40S/60S/80S peaks indicated active translation (Fig. 4C). In contrast, NT-2 treatment resulted in a pronounced accumulation of 80S monosomes, accompanied by a striking depletion of polysomes, consistent with global translation inhibition. We tested whether NT-2 indirectly inhibits translation through eIF2α phosphorylation, a central marker of the integrated stress response. No increase in phospho-eIF2α was observed after NT-2 treatment (Fig. 4D), indicating that the repression is not mediated through this signalling route.

To assess phylogenetic specificity, we extended the analysis to prokaryotic and fungal systems. *E. coli* growth was unaffected by NT-2, in contrast to tetracycline-treated controls (Fig. 4E), and NT-2 failed to inhibit protein synthesis in bacterial S30 extracts (Fig. 4F), indicating that NT-2 does not impair prokaryotic ribosomes. In *S. cerevisiae*, NT-2 similarly did not affect cellular growth (Fig. 4G). Consistently, AHA-labelling in living yeast cells showed no reduction of protein synthesis upon NT-2 treatment (Fig. 4H, Sup. Fig. 3). However, translation assays in yeast lysates revealed concentration-dependent inhibition, with ∼70% reduction at 5 µM and complete inhibition at 50 µM (Fig. 4I). These results suggest that NT-2 can inhibit both mammalian and fungal protein synthesis *in vitro*, but its growth-inhibitory effects in intact cells are species-dependent, likely reflecting differences in uptake, efflux, or metabolic modification.

### NT-2 binds the 60S PTC

To define the mechanism of NT-2 translation inhibition, we determined the structure of human 80S ribosomes from NT-2–treated HEK293T cells by single-particle cryo-electron microscopy (cryo-EM). As controls, we reconstructed human 80S ribosomes from untreated cells (CTR Ribosomes), with or without incubation in NT-2, before vitrification. Density for NT-2 was detected in the reconstructions of human 80S ribosomes from NT-2-treated HEK293T cells, and in ribosomes incubated with NT-2, but not in the untreated 80S controls, indicating that NT-2 binds ribosomes in cells and remains associated during purification (Fig. 5A-B).

**Figure 5.**
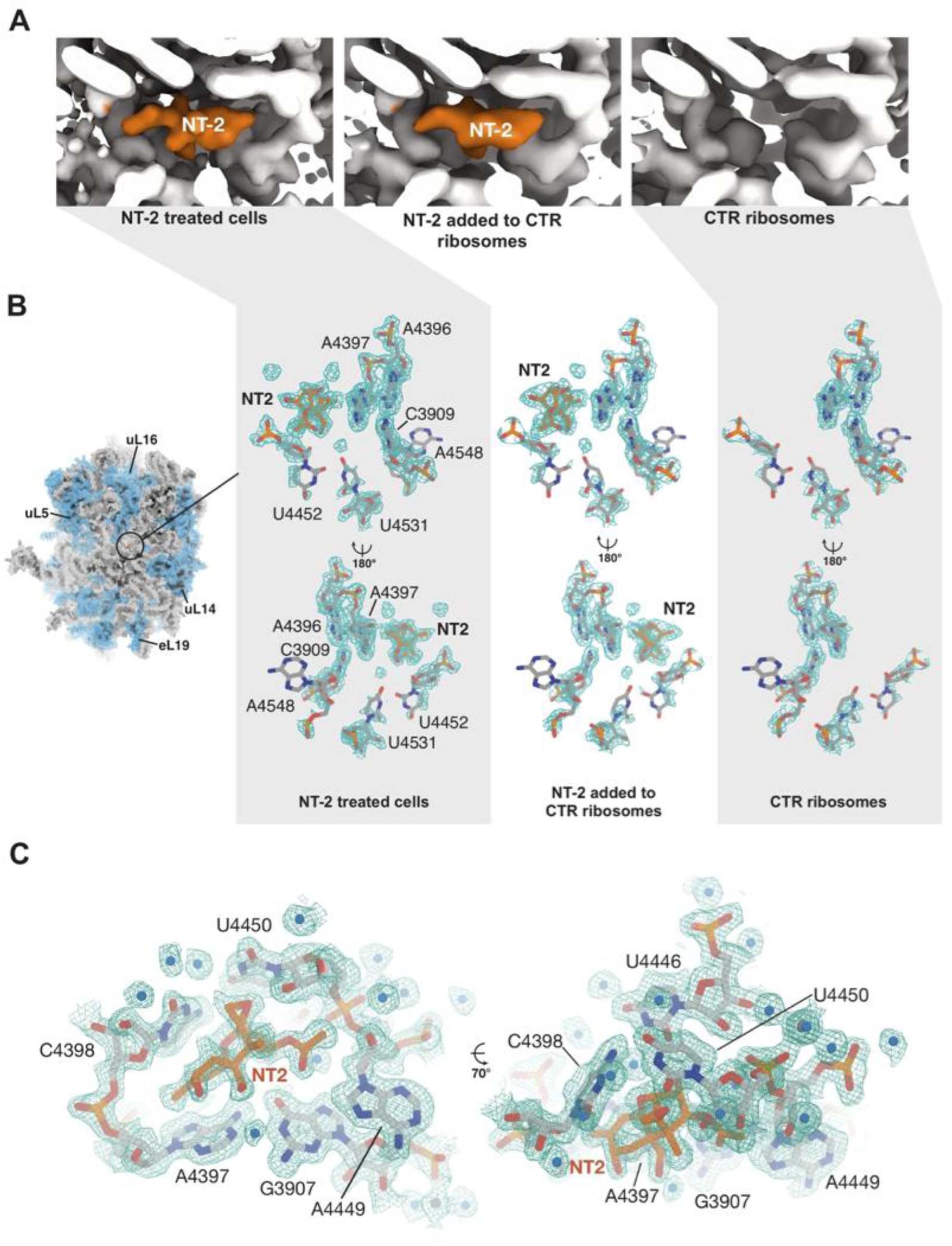
NT-2 binds the ribosomal PTC and inhibits elongation . **(A)** Cryo-EM maps of the 60S PTC from NT2-treated cells (left), NT-2 added to ribosomes from untreated cells (middle), and untreated control ribosomes (right). Clear NT-2 density (orange) is present only in the NT-2–treated maps but absent in the untreated CTR ribosomes. **(B)** Refined atomic model of the human 60S ribosomal subunit with bound NT-2 in overview (left, refined atomic structure shown in surface representation), and close-up (right, in stick representation). **(C)** NT-2 at the PTC of ribosomes of NT-2 treated cells. The refined atomic structure of NT-2 (orange) and the surrounding nucleotides (light grey) are shown in stick representation. Water molecules are shown as blue spheres, while magnesium atoms are shown as dark grey spheres. Experimental Cryo-EM density is plotted at a contour level of 4 *σ* and shown as a mesh.

The reconstruction reached a resolution of 1.76 Å, providing near-atomic detail (Fig. 5B-C, Sup. Fig. 5). The NT-2 density was clearly resolved in the PTC of the large subunit, occupying the A-site cleft. The compound showed no evidence of intracellular modification such as de-epoxidation or deacetylation. Binding is mediated by hydrogen bonds and van der Waals contacts (Fig. 5C). Key interactions include hydrogen bonds between the ethyleneoxide group of NT-2 and the C2 ribose hydroxyl group of U4450, between the C-3 hydroxyl group of NT-2 and O-2 of the uracil base U4446, and between the oxane of NT-2 and both, the cytosine base of C4398 and the uracil base of U4446. Additional van der Waals contacts involve the adenine base of A4397. The 4β-acetoxy group forms hydrogen bonds with the guanine base of G3907 and the ribose of A4449.

The lack of activity in bacterial systems is explained by divergence in the prokaryotic PTC, which cannot accommodate NT-2^31^. Together, these structural and biochemical data establish NT-2 as a eukaryote-specific inhibitor that binds the PTC .

### Molecular architecture of NT-2 binding at the ribosomal PTC

To understand further the structural context of NT-2 binding, we examined the compatibility of the compound with tRNA and nascent chain positioning within the PTC. Docking of a model P-site tRNA and nascent polypeptide chain onto the NT-2–bound ribosome indicates that NT-2 occupies the A-site cleft of the PTC in close proximity to the peptidyl-transferase active site (Fig. 6A). In this configuration, NT-2 does not sterically clash with the P-site tRNA, consistent with the observed structural accommodation of the peptidyl-tRNA within the ribosomal active site.

**Figure 6.**
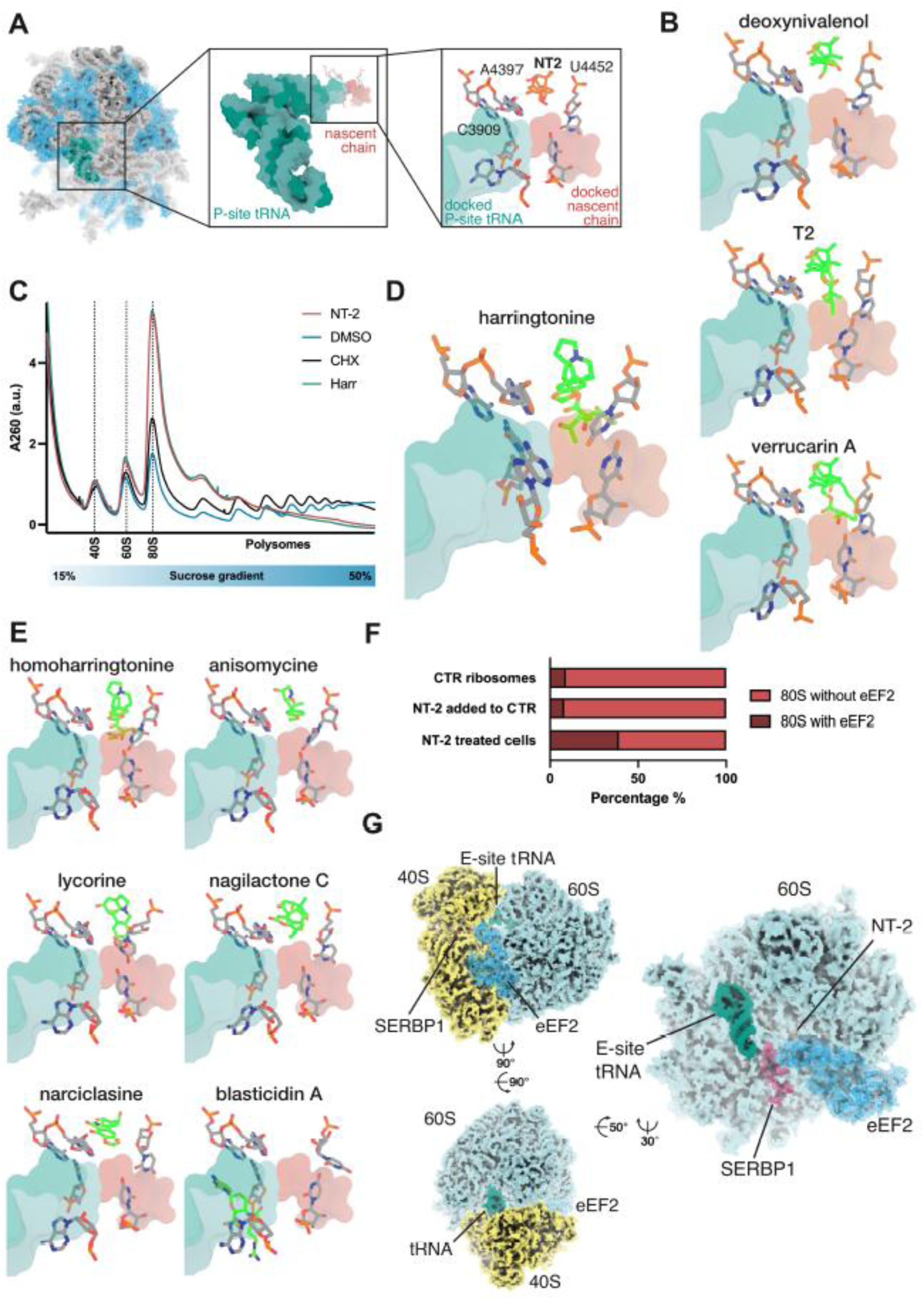
Molecular architecture of NT-2 binding and associated ribosome state. **(A)** Structural context of NT-2 binding at the PTC. The refined structure of the 60S ribosomal subunit with bound NT-2 is shown in surface representation. A tRNA bearing a nascent polypeptide chain (PDB: 7QWR; ^38^) has been modelled by rigid body docking into the active site and is shown as teal and red surface, respectively. Overlay of the tRNA and nascent chain over NT-2 suggests absence of direct sterical clashes. **(B)** Overlay of 60S cryo-EM structures of modelled trichothecene toxins deoxynivalenol, T-2 toxin and verrucarin A in the PTC. The compound and immediate environment of the PTC are shown in stick representation in magenta and light gray, respectively. The docked tRNA and nascent chain is shown as semitransparent surface. **(C)** Polysome profiles of HEK293T cells treated with NT-2 compared with DMSO, cycloheximide (CHX) and harringtonine. **(D)** Harringtonine binds in the PTC A-site cleft (PDB: 7UCJ; ^39^). Harringtonine and the immediate environment of the PTC are shown in stick representation in magenta and light gray, respectively. The docked tRNA and nascent chain is shown as semitransparent surface. **(E)** Structural comparison of NT-2 with additional PTC-targeting inhibitors including homoharringtonine, anisomycin, lycorine, nagilactone C, narcislasine and blasticidin S. **(F)** Particle distribution after classification of the entire cryo-EM particle set with focus on eEF2 indicates that ribosomes purified from NT-2 treated cells contain substantially higher numbers of eEF2-bound complexes. **(G)** Cryo-EM structure of NT-2 bound dormant ribosomes (resolution 2.08 Å). Refined atomic structure of the dormant eEF2-bound 80S state found in ribosomes purified from NT-2 treated cells. The complex contains the large (light blue surface) and small (yellow surface) ribosomal subunit, an E-site tRNA (teal surface), eEF2 (blue semi-transparent surface and cartoon), SERBP1 (magenta semitransparent surface and cartoon), and NT-2 (orange surface).

Comparison with structural data from other trichothecenes^24^ revealed a highly similar binding mode within the PTC pocket. Structural overlays of deoxynivalenol, T-2 toxin, and verrucarin A show that these compounds occupy the same region of the A-site cleft (Fig. 6B). Despite the overall scaffold similarity, subtle differences in substituents alter the spatial arrangement of individual toxins. In particular, modelling suggests that trichothecenes such as T-2 toxin and verrucarin A could interfere with the positioning of the A/A-site tRNA relative to the P-site peptidyl-tRNA, whereas NT-2 remains compatible with the P-site tRNA configuration.

Polysome profiling further revealed that NT-2 treatment results in a collapse of polysomes and accumulation of 80S ribosomes, a phenotype closely resembling that observed for harringtonine (Fig. 6C). Harringtonine is a well-characterized early elongation inhibitor that prevents productive elongation immediately after initiation^32^. Consistently, polysome gradients obtained in the presence of NT-2 closely resemble those produced by harringtonine and are clearly distinct from the profile observed with cycloheximide, which stabilizes elongating ribosomes and results in polysome arrest.

Structural comparison with harringtonine shows that both compounds occupy the same functional pocket within the A-site region of the PTC (Fig. 6D). Extending this analysis to additional PTC-targeting inhibitors, including homoharringtonine, anisomycin, lycorine, nagilactone C, narcislasine and blasticidin A ^24^, revealed that NT-2 binds within the canonical inhibitor pocket of the PTC that accommodates a wide range of structurally diverse translation inhibitors (Fig. 6E). Despite sharing this binding region, NT-2 adopts a distinct orientation relative to the surrounding rRNA nucleotides, distinguishing it from other inhibitors occupying the PTC. For example, blasticidin A binds near the P-site and interferes with peptide bond formation by stabilizing the peptidyl-tRNA in the P-site and blocking proper positioning of incoming aminoacyl-tRNA^24^.

### NT-2–bound ribosomes are associated with a dormant ribosome state

To determine how NT-2 impacts ribosome conformations, we classified cryo-EM particles of purified 80S ribosomes from DMSO-treated cells, NT-2 added to 80S and 80S from NT-2-treated cells (Sup. Fig. 6A-C). Under normal conditions, eEF2-bound rotated-2 (R2) intermediates are highly transient and rarely detected in actively translating ribosomes, whereas decoding and the A, P states are strongly represented ^33,34^. Investigation of ribosomal states by classification of subsets of the NT-2–treated sample, the DMSO control, and the NT-2 control identified an eEF2-bound 80S complex as the dominant class in the NT-2–treated sample (Fig 6F, Sup. Fig. 6E).

Classification of the complete datasets with a focus on eEF2 after subtraction of 60S density yielded three classes of eEF2-bound 80S complexes in the NT-2 sample, which all contain eEF2 and an E-site tRNA and display additional density consistent with SERBP1, and differ slightly in the conformation of the 40S subunit and eEF2 (Sup. Fig. 6D). In contrast, focused classification of the control datasets yielded one class of eEF2-bound 80S ribosomes for both the DMSO and the NT-2 control samples (Sup. Fig. 6B-C).

Quantification of ribosome states revealed that eEF2-bound 80S ribosomes constitute 39% of particles in the NT-2–treated sample, whereas only 9% and 8% of particles in the DMSO and NT-2 control samples, respectively, adopt this state (Fig. 6F). Dormant eEF2-bound ribosomes containing an E-site tRNA have been reported to represent a substantial fraction (∼20%) of ribosomes in live human cells ^35^. Consistent with this observation, we detect a smaller population of ribosomes in this configuration in purified monosomes of the control samples (8–9%). Inspection of the refined maps reveals additional density consistent with SERBP1, indicating that the eEF2-bound ribosomes observed in the control samples correspond to the dormant ribosome state.

In the NT-2–treated sample, eEF2-bound 80S ribosomes consistently display density for SERBP1 and an E-site tRNA (Fig. 6G, Sup. Fig. 6D), resulting in a ∼4-fold enrichment of this dormant configuration compared with the controls (Fig 6F, Sup. Fig. 6E). eEF2 normally engages R2 ribosomes to catalyze forward translocation, whereas SERBP1 binds translationally inactive ribosomes and stabilizes stalled ribosome–eEF2 complexes ^36^.

The simultaneous presence of eEF2 and SERBP1 therefore indicates that NT-2–bound ribosomes adopt an inactive configuration recognized by SERBP1. This configuration is distinct from canonical elongation intermediates and corresponds to an inactive, SERBP1-stabilized eEF2-bound ribosome state. Such dormant 80S ribosomes have been observed in different physiological contexts ^34,36,37^ and more recently after chemical inhibition of eukaryotic translation ^35^. Together, these cryo-EM data associate translation inhibition by NT-2 with enrichment of a SERBP1-stabilized dormant ribosome state. This state becomes prominent in cells due to eEF2 and SERBP-1 presence but is not recapitulated by the solely addition of NT-2 to purified ribosomes.

## Discussion

This study establishes a scalable, human-specific cell-free translation screening (CFTS) platform and applies it to identify novel translation modulators. Unlike cell-based assays, which often miss potent inhibitors due to cytotoxicity or poor cellular uptake, CFTS enables direct readout of protein synthesis in human lysates under controlled conditions. Using cytoplasmically enriched extracts prepared by dual centrifugation, we achieved batch consistency, freeze–thaw compatibility, and preservation of human-specific translation features, including cap-dependent initiation, regulatory RNA elements, and poly(A) sensitivity ^12,27^. Similar HeLa S3 lysates have previously been shown to recapitulate known and reveal new translation-relevant processes ^40–43^. These features, together with assay miniaturization and signal stability, support the use of CFTS as a scalable human lysate-based platform for identifying and prioritizing candidate translation modulators for further mechanistic characterization. By adapting mRNA reporters or lysate cell types ^12^, the platform can also be extended to interrogate readthrough, frameshifting, and transcript- or cell type–specific regulation.

The screen recovered both established inhibitors, including cycloheximide and puromycin, and the T-2 toxin ^2,24,30,44^, and a chemically diverse set of novel hits. (Sup. Table 3 and 4) Among these, the trichothecene NT-2 stood out as a previously uncharacterized translation modulator. First described in structural studies of Fusarium-derived mycotoxins, NT-2 exhibits an uncommon hydroxylation pattern and a 4β-acetoxy substitution, which is present in only a subset of related trichothecenes ^30^. As a sesquiterpenoid, NT-2 belongs to a natural product class related to ribotoxic and immunotoxic activities ^15,25,30^. Our data now reveal NT-2 as a natural compound that can directly and specifically inhibit eukaryotic ribosomes, confirming its cellular relevance and highlighting underestimated risks related to cereal contamination by species of the *Fusarium* genus ^16^.

The cross-kingdom profile of NT-2 highlights how physiology and ribosome evolution dictate inhibitor sensitivity. In mammalian cells, NT-2 inhibition was stronger than in lysates, consistent with enhanced uptake or potentiation by endogenous pathways, a phenomenon noted for other translation inhibitors ^45^. Conversely, NT-2 showed no detectable effect on *E. coli* growth or S30 extracts, confirming that it does not impair prokaryotic protein synthesis. This strict eukaryotic specificity implies sparing of bacterial communities such as the gut microbiota, a crucial point for toxicology and therapeutic considerations ^46,47^. In *S. cerevisiae*, NT-2 inhibited translation in lysates but not in living cells, with no impact on the yeast growth, likely due to limited intracellular access across the multilayered yeast cell wall ^48–50^. Once the barrier is bypassed, NT-2 displays potency comparable to that of other trichothecenes ^14^, consistent with a conserved eukaryotic ribosomal target such as the PTC. This divergence between *in vitro* and *in vivo* activity suggests that NT-2 is more relevant toxicologically for animal and human hosts than for suppressing fungal competitors ^13,14,16^.

At sub-2 Å resolutions, chemical modifications and solvent interactions can be directly visualized ^51,52^. Our 1.76 Å cryo-EM map of NT-2 bound to the human 60S ribosomal subunit achieves such interpretability, unambiguously placing the inhibitor in the PTC A-site cleft. This structural information is consistent with the polysome collapse with 80S accumulation observed *in vivo* (Fig. 6C), supporting interference with an early post-initiation / elongation-cycle step and the absence of eIF2α phosphorylation, indicating a direct rather than stress-mediated mechanism. The lack of inhibition in bacterial systems agrees with structural differences in the prokaryotic PTC, which lacks rRNA features contacted by NT-2 in the human ribosome ^53–55^. NT-2 is a member of the trichothecenes, a large group of toxins that share a core structure but differ in the substituents at positions 3, 4, 7, 8, and 15 (Fig. 3A) and vary in extent and mechanism of toxicity. Comparison of our structure of the human 80S with NT-2 and the structure of T-2 bound to the yeast ribosome ^24^ shows that both toxins occupy the same site in the conserved eukaryotic PTC and establish similar rRNA contacts. Larger substituents in position 3 (e.g. OAc in calonectrin) would clash with the bases of G4393 and U4446, while shorter substituents in position 4 (e.g. H in calonectrin) would abolish hydrogen bonding contacts with G3907 and A4449. Hydroxyl groups at positions 8 and 15 of NT-2 do not contact the ribosome, while the 4-Methylpent-4-enoic acid substituent of T-2 projects into the nascent protein tunnel (Fig. 6B). Thus, large substituents at positions 8 and 15 are tolerated but dispensable for binding, whereas modifications at position 3 or 4 critically influence ribosome engagement.

The accumulation of R2–eEF2 particles only in ribosomes from NT-2–treated cells, and not *in vitro*–treated ribosomes, highlights the need to study inhibitors in their physiological context. eEF2 normally binds transiently to rotated ribosomes to promote translocation ^33,34^. The additional presence of SERBP1 in these complexes indicates that these complexes are associated with a dormant state: SERBP1 occludes the mRNA channel and interacts with eEF2, locking ribosomes in an “idle” conformation incompatible with elongation ^36,37,56–58^. This expands the known role of SERBP1 as a ribosome preservation factor, previously observed in cold-treated embryos ^37^ and mirrors the yeast ortholog Stm1, which promotes dormant 80S complexes under nutrient stress via TORC1 regulation ^59,60^. The R2-SERBP1 80S signature induced by NT-2 suggest a connection between chemical inhibition and cellular pathways that preserve inactive ribosomes. A similar signature was recently reported for CHX-treated ribosomes^35^ suggesting that chemical translation inhibition may generally be associated with ribosome dormancy. We note that the initial discovery and dose-response validation were performed in HeLa S3 lysates, whereas the cellular inhibition assays, polysome profiling and cryo-EM analyses were performed in HEK293T cells. The consistent inhibition observed across these human systems supports NT-2 as a mammalian translation inhibitor, while the enrichment of the eEF2/SERBP1-bound dormant state should be interpreted in the HEK293T cellular context used for ribosome purification. Although the high-resolution structure places NT-2 within the PTC A-site cleft, we interpret NT-2 as a PTC-binding translation inhibitor that could perturb peptide-bond formation, A-site substrate accommodation, or a closely coupled early elongation step.

Together, our cell-free screening based on human lysates identified NT-2 as a potent eukaryotic translation inhibitor. The structural and polysome profiling data support a working model in which NT-2 binds the PTC A-site and perturbs eukaryotic translation at an early post-initiation or early elongation-cycle step, leading to polysome depletion and 80S accumulation. In NT-2-treated HEK293T cells, this inhibition is associated with enrichment of an inactive eEF2/SERBP1-bound 80S state (Fig. 7).The identification of NT-2 as a translation inhibitor has implications for health, agriculture, and drug discovery. *Fusarium sporotrichioides*, the fungal source of NT-2, is a major cereal pathogen, and the presence of poorly characterized trichothecenes such as NT-2 highlights risks to food safety and the global feed supply ^13^. Trichothecene-derived scaffolds could also inform the development of anticancer drugs. Pharmacological inhibition of eukaryotic translation has therapeutic potential, as illustrated by the clinically approved translation inhibitor omacetaxine mepesuccinate used for the treatment of chronic myeloid leukemia ^61^.

**Figure 7.**
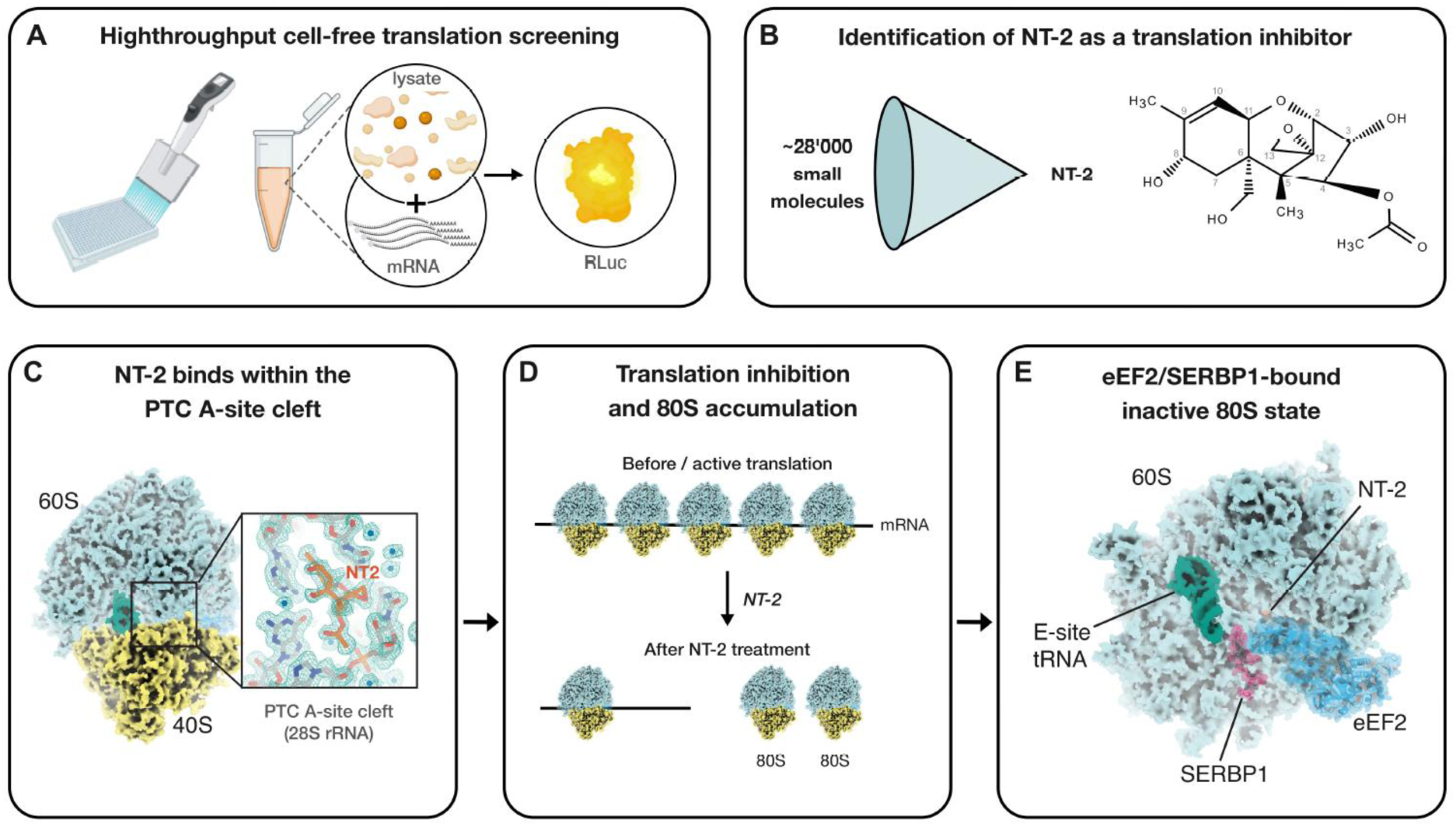
Working model for NT-2-associated inhibition of productive translation. A high throughput human cell-free translation screening identified NT-2 as a eukaryotic translation inhibitor. Cryo-EM analysis revealed that NT-2 binds within the A-site cleft of the peptidyl transferase center (PTC) of the human 60S ribosomal subunit. Polysome profiling showed polysome depletion with accumulation of 80S ribosomes, consistent with perturbation of productive translation at an early post-initiation or early elongation-cycle step. In NT-2-treated HEK293T cells, ribosomes were enriched in a dormant eEF2/SERBP1-bound 80S state. The dashed arrow indicates an associated cellular ribosome state rather than direct causality.

Overall, this work characterizes NT-2 as a potent, eukaryote-specific ribosome inhibitor that binds the PTC and is associated with SERBP1-containing dormant ribosomes. More broadly, our scalable human cell-free translation screening platform provides a high-resolution strategy for discovering translation-modulating compounds and linking environmental toxins to ribosome biology.

## METHODS

### Cell lines and cell culture

HeLa S3 cells (Sigma-Aldrich, ECACC, Cat. No. 87110901) and HEK293 Flp-In T-REx cells (Invitrogen Cat. No. R78007) were cultured in DMEM/F-12 (Gibco, Cat. No. 32500–043) containing 10% FCS (SIGMA, Cat. No. F7524-500ML), 100 U/ml Penicillin-Streptomycin (Bio-Concept, Cat. No. 4-01F00-H) (termed DMEM+/+) at 37°C and 5% CO_2_ in humid atmosphere. HEK293T cells were detached with Trypsin-EDTA PBS (BioConcept, Cat. No. 5-51F00-H). Cell counting was performed with Trypan Blue Stain (Gibco, Cat. No. 15250-061) using the automated LUNA II cell counter (Logos Biosystems, Cat. No. L40002).

### Preparation of HeLa translation-competent lysate

Lysates were prepared from HeLa S3 cell cultures like previously ^27^ grown to a density ranging from 1 to 2 x 10^6^ cells/ml, pelleted at 200 x g, 4°C for 5 min, washed twice with cold 1x PBS (pH 7.4) and resuspended in a translation-competent buffer, (33.78 mM HEPES (pH 7.3), 63.06 mM KOAc, 0.68 mM MgCl2, 54.05 mM KCl, 13.51 mM creatine phosphate, 229.73 ng/µL creatine kinase and 1x protease inhibitor cocktail (Bimake, Cat. No. B14002)) to a final concentration of 2 x 10^8^ cells/ml in 2 mL screw cap microtubes (Sarstedt, Cat. No. 72.693). The cells were lysed by dual centrifugation at 500 RPM at -5°C for 4 min using the ZentriMix 380 R system (Hettich AG) with a 3206 rotor with 3209 adapters. After dual centrifugation, the lysate was pelleted at 13’000 x g, 4°C for 10 min. The supernatant (the lysate) was aliquoted, snap-frozen and stored at -80°C. The lysate could be thawed and refrozen multiple times for subsequent applications with only a minor loss of translation efficiency as previously reported^27^.

### Preparation of reporter RLuc mRNA

Reporter mRNAs were encoded from plasmids with a pCRII backbone. The 3xFLAG-RLuc (humanized) reporter contains an AG initiator sequence for capping with the CleanCap reagent AG, the human beta globin 5’ UTR, a 20 bp 3’ UTR, a 30 nucleotide-long template-encoded poly(A) sequence and a BsmFI binding site downstream for linearization. The previously described ^43^ p200-RLuc (humanized) reporter contains a 70 bp 5’UTR, a 500 bp 3’ UTR with 6xMS2, an 80 nucleotide-long template-encoded poly(A) sequence and a *Hind*III binding site downstream for linearization.

Before *in vitro* transcription, the plasmids were linearized using *BsmF*I (New England Biolabs, Cat. No. R0572 L) or *Hind*III-HF (New England Biolabs, Cat. No. R3104L) in reactions containing 40 ng/ml DNA (8 µg in total) in 1x CutSmart buffer (New England Biolabs, Cat No. B6004S) overnight at 37°C. After confirming linearization by agarose gel electrophoresis, the linearized DNA was purified and concentrated using the ChIP DNA Clean & Concentrator kit (ZYMO research, Cat No. D5205) and eluted in 10 µl elution buffer. *In vitro* transcription, using the linearized plasmid as a template (10 µl of elution), was performed in a 100 µL reaction containing 1x T7 transcription buffer (Thermo Fisher Scientific, Cat No. EP0111), 1 mM of each NTP (Thermo Fisher Scientific, Cat No. R0481), 1 U/ml RNase inhibitor (Vazyme, Cat No. R301–03), 0.001 U/ml pyrophosphatase (Thermo Fisher Scientific, Cat No. EF0221) and 1.5 U/ml T7 polymerase (Thermo Fisher Scientific, Cat No. EP0111). For the 3xFLAG-RLuc reporters, 0.8 mM CleanCap reagent AG (TriLink Biotechnologies, Cat No. N-7113–5) was included in the reaction. The transcription reaction was incubated at 37°C 1 h. An additional 1.5 U/ml T7 polymerase was added, and the reaction was further incubated for 1 h at 37°C. Subsequently, 0.15 U/ml Turbo DNase (Invitrogen, Cat No. AM2238) was added, and the reaction was incubated at 37°C for 30 min to digest the plasmid DNA. The transcribed mRNA was isolated using the Monarch RNA Cleanup Kit (New England Biolabs, Cat No. T2040 L), eluted in 1 mM sodium citrate pH 6.5 (Invitrogen, Cat. No. AM7001), and quantified by A260 measurement.

The p200-RLuc reporter was capped using the Vaccinia Capping System (NEB, Cat. No. M2080S) according to the manufacturer’s instructions, with the modification that 1 U/ml RNase inhibitor (Vazyme, Cat. No. R301–03) was added to the reaction. The capped mRNA was isolated and quantified as described above after transcription. All *in vitro* transcribed mRNAs were aliquoted, snap-frozen, and stored at -80°C.

### High throughput screening

Compounds from the screening library, along with DMSO and cycloheximide controls, were pre-spotted into 384-well plates (Greiner Bio-One CELLSTAR, Cat. No. 781073) at 5 nL, 10 mM using the LABCYTE Echo 655 and Access Workstation. Plates were stored at -20 °C until use in the BSF Screening facility at the EPFL. A translation-competent premix was prepared in advance by diluting HeLa S3 lysate to a concentration of 1 × 10^8^ cell equivalents/ml, adding RLuc reporter mRNA at a final concentration of 5 fmol/µL and Murine RNase inhibitor (Vazyme, Cat. No. R301-03-AA) at a final concentration of 1 U/µL, frozen in liquid nitrogen and transferred to the BSF facility in dry ice. Using the BioTek MultiFlo dispenser, 5 μL of lysate was added to each well, yielding a final compound concentration of 10 µM. Translation reactions were incubated at 33°C for 50 minutes. Subsequently, 5 μL of 1x Renilla-Glo substrate (Promega, Cat. No. E2720) was added, followed by a 10-minute incubation at room temperature. Luminescence was measured using the BioTek Synergy Neo Plate Reader. The Z′ factor was calculated as described ^29^, measuring the separation between positive and negative controls.

### Hela S3 cell-free translation assays

For *in vitro* translation, the HeLa S3 lysate was used at a concentration of 1 × 10^8^ cell equivalents/ml (stock = 2 × 10^8^ cell equivalents/ml) and supplemented with 1 U/ml RNase inhibitor (Vazyme, Cat. No. R301-01-AA). The *in vitro* transcribed and capped mRNAs were incubated at 65°C for 5 min and cooled on ice before addition. Unless otherwise stated, *In vitro* translation was performed at 37°C for 1 hour in 12.5 µL reactions and stopped on ice. To evaluate the effect of NT-2 on translation, the following conditions were tested in parallel: water as negative control for inhibition (CTR), NT-2 at 0.05 µM, 0.5 µM, 1.5 µM, 3 µM, 5 µM, 15 µM 50 µM and 100 µM, or 1 mg/ml puromycin (Santa Cruz Biotechnology, Cat. No. sc-108071) as positive control for translation inhibition.

To monitor the protein synthesis output by the luciferase assay, 12.5 µL of each translation reaction was mixed with 12.5 µL 1x Renilla-Glo substrate in Renilla-Glo assay buffer (Promega Cat. No. E2720) and supplemented with 35 µL of H_2_O in a white-bottom 96-well plate (Greiner, Cat. No. 655073). The luminescence was measured with the Promega GloMax Explorer GM3500 plate reader after 10 min of incubation at room temperature with 0.3s integration time per well. Three luminescence measurements were taken from each biological replicate, which were averaged and plotted.

### Polysome profiling

75–90% confluent HEK293 cells were cultured in 150 mm dishes and treated with either 2 μM NT-2 Toxin, 100 µg/ml cycloheximide (CHX), 2 µg/ml harringtonine, or 0.02% DMSO (CTR) in DMEM+/+ medium for 2 h at 37 °C and 5% CO_2_. After medium removal, the cells were scraped in ice-cold PBS and transferred to a 2 ml microcentrifuge tube. The cells were pelleted for 5 min at 4°C and 500 x g, washed with ice-cold PBS, pelleted again and resuspended in 1x lysis buffer (10 mM HEPES pH 7.5 (Thermo Scientific Cat. No. J61275.AK), 100 mM KCl (Thermo scientific, Cat. No. J62422.AK), 5 mM MgCl_2_ (Jena Bioscience, Cat. No. BU-110-1M), 1 mM Triton X-100 (Thermo Scientific, Cat. No. 327371000), 1% Sodium deoxycholate (SIGMA-ALDRICH, Cat. No. D6750-25G), 1 mM DTT (Merk, Cat. No. 43816-10mL), 0.1 U/µL Murine RNase inhibitor (Vayzme, Cat. No. R301-01-AA), 0.1 x Protease inhibitor cocktail (Bimake, Cat. No. B14002) and 0.0125 U/µL Turbo DNase (Invitrogen, Cat No. AM2238) and incubated for 2 min on ice. After incubation on ice, the cell lysate was centrifuged for 5 min at 4°C and 16,000 x g, and the supernatant was collected.

15% and 50% (w/v) sucrose solutions (20 mM HEPES pH 7.5, 200 mM KCl, 10 mM MgCl_2_ in H_2_O) were supplemented with 1 mM DTT (Merck, Cat. No. 43816-10mL) and gradients were generated using a Biocomp Gradient Station ip in SW41 tubes (SETON, Cat. No. 7030).

300 µL of lysed cells were loaded onto the gradients and the tubes were spun in the SW 41 Ti rotor for 2 h at 274355 x g, 4°C in a BECKMAN COULTER Optima XPN-80 ultracentrifuge. The gradients were fractionated using the Biocomp PGF ip Piston Gradient Fractionator at a speed of 0.3 mm/s in 20 microcentrifuge tubes per replicate. During fractionation, the A260 profile was recorded, and absorption measurements were visualized using Graph Pad Prism (version 10.0.2). The sucrose fractions were directly used for Cryo-EM sample preparation.

### Cryo-EM sample preparation

For cryo-EM, sucrose was removed from the collected and combined monosome fraction using the RAPPL protocol ^62^. Briefly, the selected fraction from the NT-2 treated sample and the DMSO treated control was diluted 1:5 with wash buffer (100 mM HEPES pH 7.5, 50 mM KCl, 10 mM Mg(OAc)_2_, 1 mM DTT, 1x protease inhibitor cocktail and 0.1 U/µL Murine RNase inhibitor (Vayzme, Cat. No. R301-01-AA). Poly-lysine magnetic beads (MCLAB, Cat. No. PLYSMB-50) were equilibrated with wash buffer, then combined with the diluted sample and incubated for 30 min at 4 °C under rotation. The beads were separated using a magnetic rack, washed three times with wash buffer and eluted in 50 µL elution buffer (wash buffer + 2 mg/mL poly-glutamic acid (SIGMA-ALDRICH, Cat. No. P4761-25MG)) for 30 min at 4 °C on a thermomixer (1,000 rpm). The eluate was quantified by A_260_ measurement and diluted with elution buffer as needed to reach an absorbance between 4 and 8 A_260_ units. Samples were snap-frozen in liquid nitrogen and stored at -80 °C until further use.

### Vitrification

Single particle cryo-EM samples were vitrfied using the plunge freeze technique ^63^. Amorphous carbon with a thickness of 1.0 nm was deposited by evaporation onto freshly cleaved mica in a CCU-010 carbon coater (Safematic, Zizers, Switzerland). Continuous carbon foil was deposited on R2/2 holey carbon grids (Quantifoil, Großlöbichau, Germany) by floatation in distilled water. The grids were subjected to negative glow discharge in a PELCO easiGlow glow discharger (Ted Pella, Redding CA, USA) at 15 mA for 15 s prior to vitrification. The climate chamber of a Vitrobot Mark 4 (Thermo Fisher Scientific, Waltham MA, USA) was equilibrated to 4°C and the humidity was adjusted to 95%. 4 µl of protein sample were applied to the grid and the sample was incubated on the grid for 30 s. The grid was subsequently blotted for 2.5 s and - without further incubation - plunged immediately into liquid ethane-propane mixture maintained at liquid nitrogen temperature. The grid was stored submerged in liquid nitrogen until data collection.

### Cryo-EM data collection

Single-particle cryo-EM data was collected on a Titan Krios G3i cryo-electron microscope (Thermo Fisher Scientific, Waltham MA, USA) equipped with a GIF BioQuantum energy filter and a K3 direct electron detector (Gatan, Pleasanton CA, USA). Micrographs were recorded in superresolution counting mode, using a magnification of 130,000 x, resulting in a pixel size of 0.648 Å/pixel. The energy filter was set to a slit width of 20 eV. Micrographs for both datasets were acquired with a total dose of 50 e^−^/Å^2^ (NT-2 incubated sample) and 60.3 e^−^/Å^2^ (controls). Automated image acquisition with aberration-free image shift was controlled by EPU software (Thermo Fisher Scientific, Waltham MA, USA). Defocus was adjusted between -0.8 µm and - 2.8 µm in 0.2 µm increments.

### Cryo-EM image processing, model building, and refinement

Image processing was carried out in cryoSPARC (Structura Biotechnology, Toronto, Canada) as outlined in the scheme Sup. Fig. 4 and 6. The atomic model of the 60S ribosomal subunit of human ribosome (PMID: 39658079) was rigid body fitted into the cryo-EM density map in UCSF Chimera (PMID: 15264254). The model was rebuilt in coot (PMID: 20383002) and refined using phenix real space refinement (PMID: 31588918).

### Puromycin incorporation & eIF2 alpha assay

HEK293 or BHK-21 cells were seeded in DMEM+/+ 24h before treatment and grown to approximately 80% confluency. For the puromycin incorporation assay, cells were treated with DMEM +/+ containing different concentrations of NT-2 (2 µM, 1 µM, 0.5 µM, 0.25 µM, 0.125 µM, 0.0625 µM, 0.03125 µM), 0.02% DMSO as a vehicle control (matching the concentration in the highest NT-2 dilution), or cycloheximide (100 µg/mL) as a positive control for translation inhibition. After 90 minutes of treatment, puromycin (10 µg/mL) was added for 10 minutes, followed by a 30-minute recovery in the respective treatment medium. For the eIF2α phosphorylation assay, cells were treated with 2 µM NT-2 or 0.02 % DMSO control for 2h.

After treatments, cells were washed with PBS, detached using trypsin, and collected by centrifugation. Pellets were washed twice with cold PBS, then lysed in RIPA buffer (50 mM Tris-HCl pH 8.0, 150 mM NaCl, 1% NP-40, 0.5% sodium deoxycholate, 1% SDS) containing 1x Protease inhibitor cocktail (Bimake, Cat. No. B14002); for the eIF2α phosphorylation assay 1x phosphatase inhibitor cocktail (Biotool, Cat. No. B15001) was also included. Samples were incubated on ice for 20 minutes, with occasional gentle mixing by flicking. The lysate was cleared by centrifugation and stored at -80 °C until Western blot analysis.

### Western blot analysis

For Western blot analysis of *in vitro* translation reactions, 4 µL of each reaction (described above) was loaded onto a mPAGE 4–12% Bis-Tris 12-well gel (Millipore, Cat. No. MP41G12). For cell lysates from puromycin incorporation and eIF2α assay, volumes were adjusted to correspond to 1 µg of total RNA per lane, and samples were loaded onto a mPAGE 4–12% Bis-Tris 10-well gel (Millipore, Cat. No. MP41G10). Gels were run for 45 minutes in MES running buffer (50 mM MES (SIGMA, Cat. No. M3671-250G), 50 mM Tris base (Fisher Bioreagents, Cat. No. BP152.5), 0.1% SDS (Fisher Bioreagents, Cat. No. BP2436-1), 1 mM EDTA (SIGMA-ALDRICH, Cat. No. E5134-500G), pH 7.3).

Proteins were transferred onto nitrocellulose membranes using the Trans-Blot Turbo Transfer Pack (BIO-RAD, Cat. No. 1704158). Membranes were then cut according to molecular weight of the target proteins and blocked for 30 minutes in 5% milk in TBS-Tween (0.1%). Primary antibody incubations were performed overnight at 4 °C under gentle agitation using the following antibodies:

Mouse anti-FLAG M2 (Sigma, Cat. No. F1804, 1:1000),

Mouse anti-Vinculin (Santa Cruz Biotechnologies, Cat. No. sc-73614, 1:2000),

Mouse anti-Puromycin (Millipore, Cat. No. MABE343, 1:15’000, IgG2a),

Mouse anti-GAPDH (Santa Cruz Biotechnology, Cat. No. sc-47724, 1:10’000, IgG1),

Mouse anti-eIF2α (Cell Signaling Technology, Cat. No. 2103, 1:1000),

Rabbit anti-phospho-eIF2α (Ser51) (Cell Signaling Technology, Cat. No. 9721, 1:1000).

After washing twice for 10 minutes in TBS-Tween (0.1%), membranes were incubated with fluorescent secondary antibodies:

Donkey anti-mouse 680LT (LI-COR, Cat. No. 926–68022, 1:10’000),

Donkey anti-rabbit 800CW (LI-COR, Cat. No. 926–32212, 1:10’000),

Goat anti-mouse IgG2a 800CW (LI-COR, Cat. No. 926-32351, 1:10’000),

Goat anti-mouse IgG1 680LT (LI-COR, Cat. No. 926-68050, 1:10’000),

Goat anti-mouse 680LT (LI-COR, Cat. No. 926-68020, 1:10’000).

Secondary antibody incubations were carried out for 1 hour at room temperature under gentle agitation. Membranes were washed twice for 10 minutes in TBS-Tween (0.1%) before signal detection using the Odyssey Infrared Imaging System (LI-COR). For *in vivo* IC_50_ determination, the signal intensities of the puromycin and GAPDH bands were quantified using ImageJ (NIH). Puromycin signals were normalized to GAPDH to account for loading differences, and relative translation levels were calculated accordingly.

### *E. coli* cell-free translation

*In vitro* translation reactions were performed using the S30 Extract System for Linear Templates (Promega, Cat. No. L1030) according to the manufacturer’s instructions. Briefly, translation reactions (total volume: 25 µL) were assembled by mixing S30 Premix, S30 extract, and complete amino acids mix at the recommended concentrations. Additionally, the reactions were supplemented with 1U/µL Murine RNase inhibitor (Vazyme, Cat. No. R301-01-AA). *In vitro* transcribed, uncapped RLuc reporter mRNA was added at a concentration of 125 fmol/µL. To evaluate the effect of NT-2 on bacterial translation, the following conditions were tested in parallel: addition of water (baseline), NT-2 at 5 µM and 50 µM, DMSO (0.5%) as vehicle control, tetracycline as a positive control for translation inhibition (10 ng/µL), and cycloheximide (CHX, 100 µg/mL).

Reactions were incubated at 37 °C for 2 hours, and translation products were analyzed by luciferase assay as described above, by mixing 25µL of each reaction with 25 µL 1x Renilla-Glo. Negative controls without RNA were included to assess background translation levels.

### Bacteria growth curve

The effect of NT-2 on *E. coli* growth was evaluated using the BL21 (DE3) pLysS strain. An overnight culture was diluted in fresh LB medium to an initial optical density (OD_600_) of 0.05, and 400 µL aliquots were dispensed into an untreated 48-well plate (Greiner Bio-One, Cat. No. 677180). The following conditions were tested in triplicate: no additive (untreated control), DMSO (0.08%) as vehicle control, NT-2 (2 µM), tetracycline (10 µg/mL) as a positive control for growth inhibition, and LB medium only as a background (no bacteria) control.

The culture plates were incubated at 37 °C with continuous shaking, and optical density at 600 nm (OD_600_) was measured every 15 minutes over an 8-hour period using a Tecan Infinite M1000 plate reader. Growth curves were analyzed to assess the effect of each treatment on bacterial proliferation.

### Yeast cell-free translation

For the preparation of translation-competent lysates from *S. cerevisiae*, we followed the protocol described in ^64^ with some adaptations. Yeast cells (PJ69-4a strain) were grown aerobically in YPD medium. A starter culture was first grown at 30 °C with agitation until the stationary phase and subsequently used to inoculate large-scale cultures. Cells were cultivated overnight in YPD at 30 °C with vigorous shaking until reaching a density of approximately 1–2 x 10^7^ cells/mL (OD_600_ ≈ 1). Cells were harvested by centrifugation at 500 x g at 4 °C for 10 min, washed once with cold sterile water and once with 1 M sorbitol.

Spheroplasts were generated by incubation of the washed cell pellet in 10 ml spheroplasting buffer (1 M sorbitol, 10 mM Tris-HCl pH 7.5) per g of wet pellet supplemented with 1 mg/ml zymolyase (20T, Carl Roth, Cat. No. 9324.1) and 1mM DTT (Merk, Cat. No. 43816-10mL). The suspension was gently agitated at 30 °C in the dark for 1.5h, while cell wall digestion was assessed by a decrease in OD_800_. Spheroplasts were collected by centrifugation at 700 x g at 4 °C for 15 min, washed twice with cold spheroplasting buffer, and resuspended in 0.5 ml of ice-cold translation buffer (100 mM HEPES pH 7.3, 2.5 mM Mg-acetate, 120 mM K-acetate, 100 mM sucrose, 1 µM spermidine, 2.5 mM ATP, 1 mM GTP, 1 mM DTT, 20 mM creatine phosphate, 200 µg/ml creatine kinase and 1x protease inhibitor cocktail) per g of original wet pellet. Cells were disrupted on ice using a glass Dounce homogenizer. The homogenate was cleared by centrifugation at 16,000 × g for 10 min at 4 °C, and the supernatant was carefully collected while avoiding both the lipid layer and pellet. The resulting extracts were snap-frozen in liquid nitrogen and stored at -80 °C.

*In vitro* translation reactions were set up in a final volume of 12.5 µL containing: 8.75 µL yeast cell-free lysate, 1 U/µL Murine RNase inhibitor, amino acid mixture (0.04 mM of each AA), 5 fmol/µL capped and polyadenylated *in vitro* transcribed mRNA (3x-FLAG RLuc) and RNase free water. *In vitro* translation was performed at 30°C for 1 hour and stopped on ice. To evaluate the effect of NT-2 on fungal translation, the following conditions were tested in parallel: no supplement (CTR), NT-2 at 5 µM and 50 µM, DMSO (0.5%) as vehicle control, or 1 mg/ml puromycin (Santa Cruz Biotechnology, Cat. No. sc-108071) as positive control for translation inhibition. To monitor the protein synthesis output by the luciferase assay, each translation reaction was mixed with 40 µL 1x Renilla-Glo substrate in Renilla-Glo assay buffer (Promega Cat. No. E2720) and transferred into a white-bottom 96-well plate (Greiner, Cat. No. 655073). The luminescence was measured with the Promega GloMax Explorer GM3500 plate reader after 10 min of incubation at room temperature with 0.3s integration time per well. Three luminescence measurements were taken from each biological replicate, which were averaged and plotted.

### Yeast growth curve

The effect of NT-2 on *S. cerevisiae* growth was assessed using the PJ69-4a strain. An overnight culture grown in YPD medium was diluted to an initial optical density (OD_600_) of approximately 0.1, and 400 µL of the diluted culture was transferred into each well of an untreated 48-well plate (Greiner Bio-One, Cat. No. 677180). The following conditions were tested in triplicate: no additive (untreated control), DMSO (0.25%) as vehicle control, NT-2 (25 µM), cycloheximide (CHX, 10 µg/mL) as a positive control for growth inhibition, and YPD medium only (no cells) as a background control.

Plates were incubated at 30 °C with continuous shaking, and OD_600_ measurements were recorded every 15 minutes for 19 hours using a Tecan Infinite M1000 plate reader. Growth kinetics were analyzed to compare the impact of the different treatments on yeast proliferation.

### Yeast AHA labelling

To assess the effect of NT-2 on the translation rate in *S. cerevisiae,* PJ69-4a yeast cells were grown and labeled with L-azidohomoalanine (AHA) following the protocol described in ^65^ with minor modifications. Briefly, a single colony was inoculated in YPD and grown overnight at 30 °C. The next day, cultures were diluted to OD600 ≈ 0.1 in fresh synthetic complete medium and grown to mid-log phase (OD600 ≈ 0.4). Cells were collected, washed with sterile water, and resuspended in SC medium lacking methionine but supplemented with L-AHA (TargetMol, Cat. No. T41090). Treatments were applied for the pulse-labelling step as follows: 25 µM NT-2, 0.25 % DMSO (vehicle control), or 50 µg/mL cycloheximide (CHX). An additional control was incubated in YPD medium to assess background fluorescence of non-labelled yeast. After incubation, cells were permeabilized and subjected to click chemistry with an alkyne-fluorophore (AAT Bioquest, Cat. No. 1700) to label newly synthesized proteins. Fluorescence was measured by flow cytometry, and mean fluorescence values were used to quantify relative translation rates under each condition.

## Data availability

The data that support the findings of this study are available from the corresponding author upon reasonable request.

## Code availability

Does not apply

## Acknowledgments

This work was supported by Grants awarded to EDK from the Swiss National Science Foundation (SNSF CRSK-3_220624), the Multidisciplinary Center of Infectious Diseases from the University of Bern (MCID), the Fondation Claude et Giuliana, the Forschungsstiftung of the University of Bern, and the Holcim Stiftung. Figure 1A was made using Biorender. The screening was performed at the Biomolecular Screening Core Facility, EPFL, Lausanne, Switzerland, and the Cryo-EM analysis was conducted at the ETH Zurich Cryo-EM Knowledge Hub, Zurich, Switzerland, in collaboration with the NCCR RNA and Disease. We thank Wojciech Teodorowicz and Nikolaos Kouvelas for their critical reading of the manuscript.

## Author contributions

**N.S., D.A., J.L., J.R.:** Data curation, Formal analysis, Investigation, Methodology, Validation, Visualization, **M.C., G.T.:** Methodology, Project administration, Supervision, Validation, Visualization **E.K.:** Funding acquisition, Project administration, Supervision, Writing –original draft **All authors:** Writing – review & editing

## Supplementary Material

**Supplementary Figure 1.**
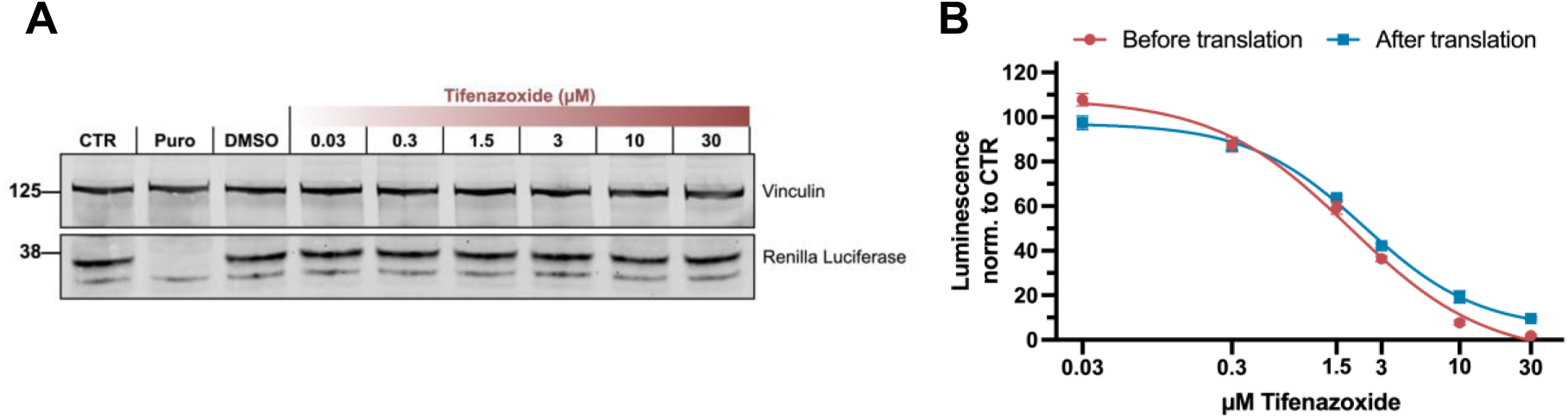
Characterization of Tifenazoxide as a *Renilla* luciferase inhibitor. **(A)** Western blot of an *in vitro* translation assay supplemented with tifenazoxide. Cell-free translation reactions were performed with *Renilla* luciferase mRNA in HeLa S3 lysate with increasing Tifenazoxide concentrations (30 nM - 30 μM) and were analyzed by SDS-PAGE and immunoblotting for *Renilla* luciferase (translation product) and Vinculin (loading control). **(B)** Cell-free translation reactions as described in (A) were performed with Tifenazoxide supplementation either before (red) or after (blue) translation. The measured luminescence was normalized to the negative control containing no supplement (CTR). Data are presented as mean values of three biological replicates (sets of translation reactions) averaged after three measurements ± SD.

**Supplementary Figure 2.**
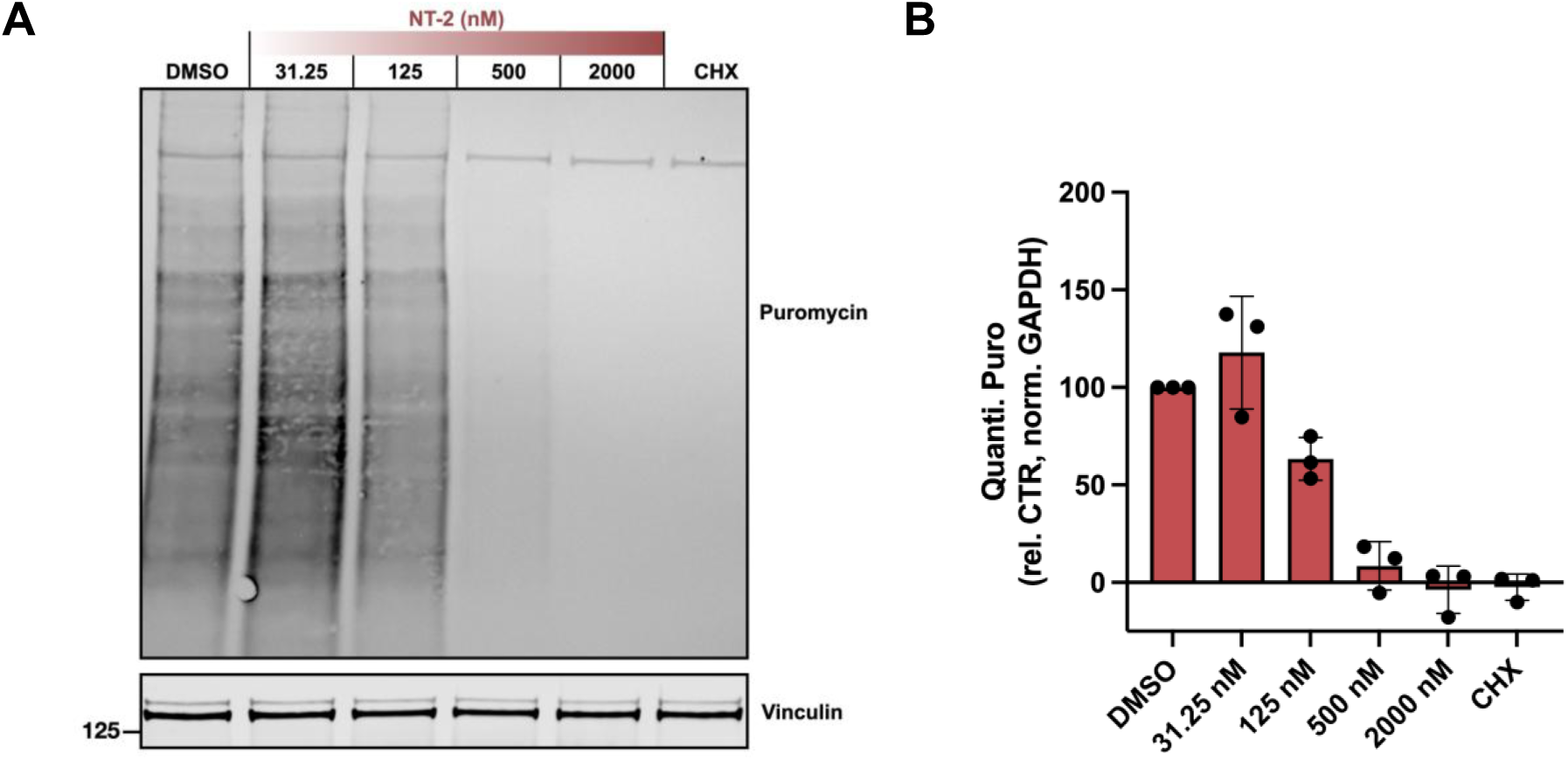
Effect of NT-2 on translation rates in *BHK-21 cells*. **(A)** Puromycin incorporation assay in BHK-21 cells treated with increasing NT-2 concentrations, DMSO, or CHX. Equal amounts of cell lysate were loaded into a 4-12% Bis-Tris gel and analysed by Western blot probing with anti-puromycin and anti-GAPDH antibodies. **(B)** Quantification of protein synthesis based on signal intensity in the anti-Puromycin-stained blot (Sup Fig 3A) normalized with GAPDH. Bars represent average of three biological replicates (dots) ± SD.

**Supplementary Figure 3.**
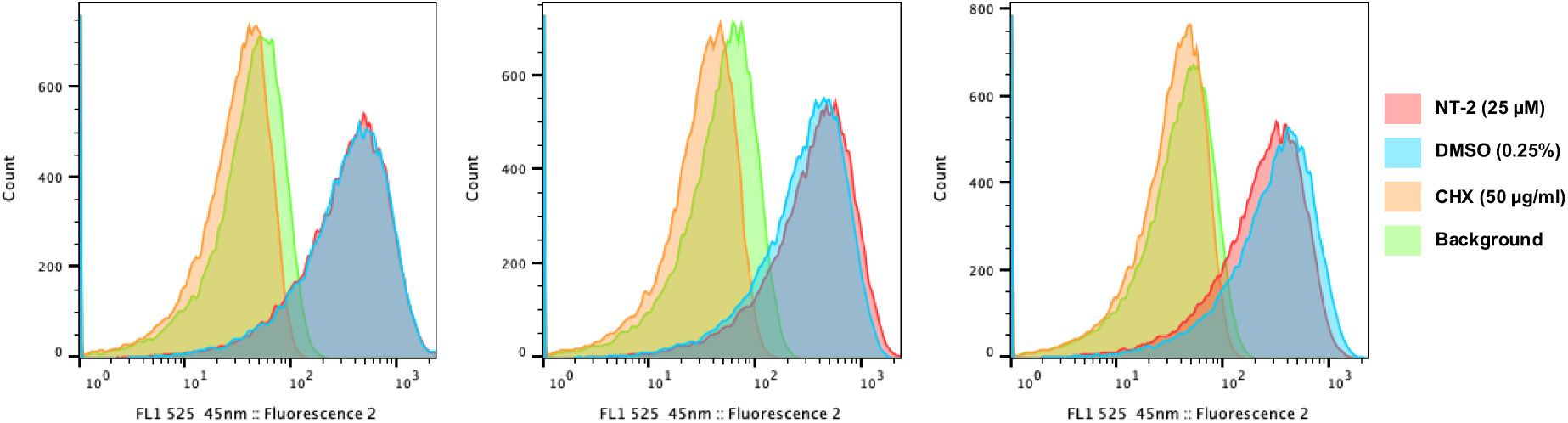
Effect of NT-2 on translation rates in *S. cerevisiae*. Fluorescence intensity of AHA-labeled yeast cells under different treatments. Cells were treated with NT-2 (25 µM), DMSO (0.25 %, control), CHX (50 µg/mL), or incubated in complete YPD medium (background control) and processed as described in Materials & Methods. Histograms show the distribution of single-cell fluorescence measured by flow cytometry of the three biological replicates.

**Supplementary Figure 4.**
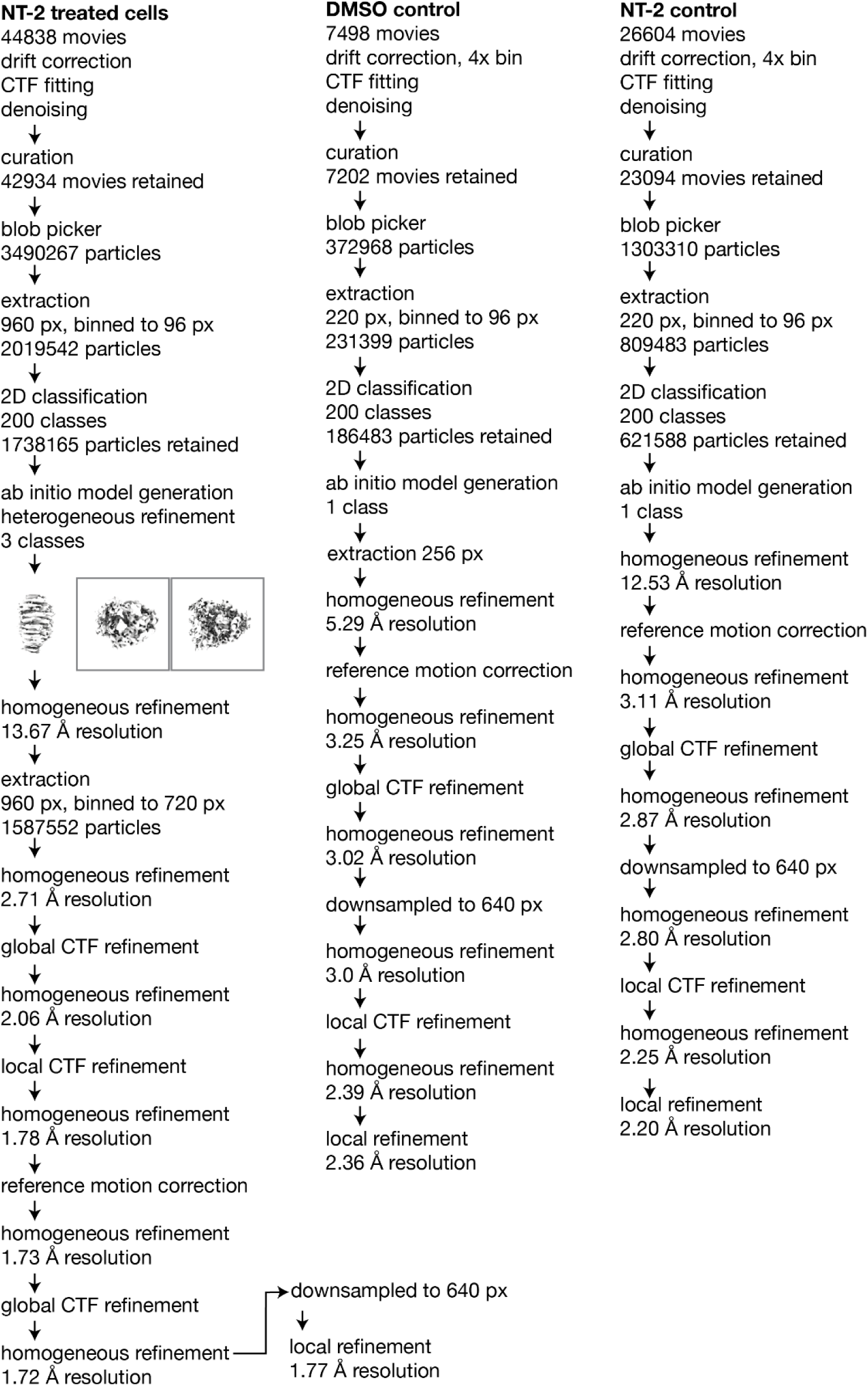
Cryo-EM data processing flowcharts. Image processing workflows for the three datasets: 80S ribosomes isolated from NT-2–treated cells, 80S isolated from DMSO-treated (CTR) cells and NT-2–incubated 80S ribosomes. Flowcharts summarize particle selection, classification, and refinement steps leading to the final reconstructions.

**Supplementary Figure 5.**
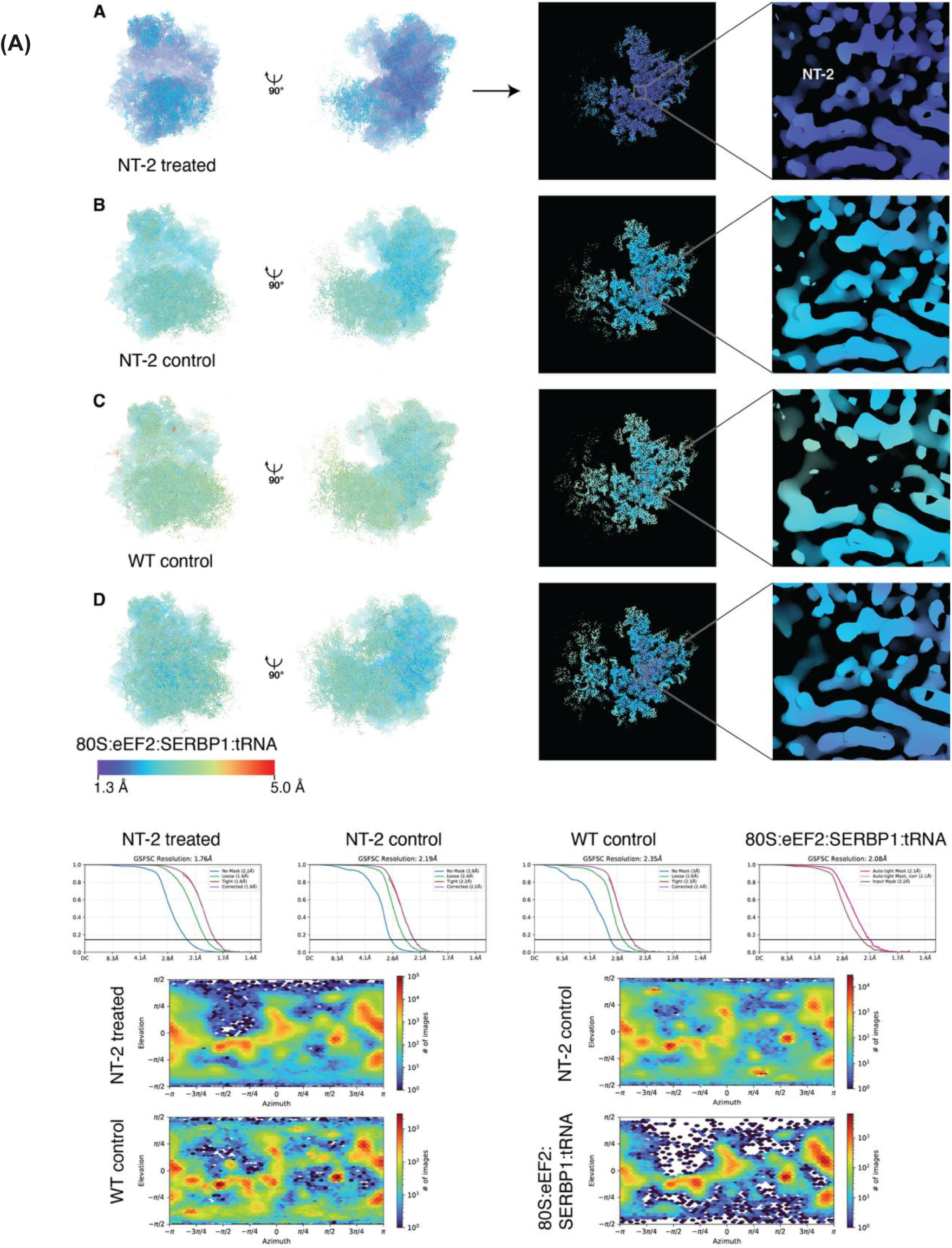
Resolution maps, FSC curves and particle orientation distribution plots. Resolution map of monosomes from NT-2 treated cells, focused on the 60S subunit. A resolution map of the surface of the molecule is shown on the left two panels. The center right panel shows a section through the map around the binding site of NT-2. On the right panel, NT-2 and the region directly adjacent are shown. All resolution maps are shown on the same scale (see inset below panel D). **(B)** Resolution map of purified monosomes incubated with NT-2 before vitrification, focused on the 60S subunit. **(C)** Resolution map of purified monosomes, focused on the 60S subunit. NT-2 is absent from the binding site. **(D)** Resolution map of dormant 80S:eEF2:SERBP1:tRNA complex purified from NT-2-treated cells. **(E)** Fourier shell correlation curves. F) Particle distribution plots.

**Supplementary Figure 6.**
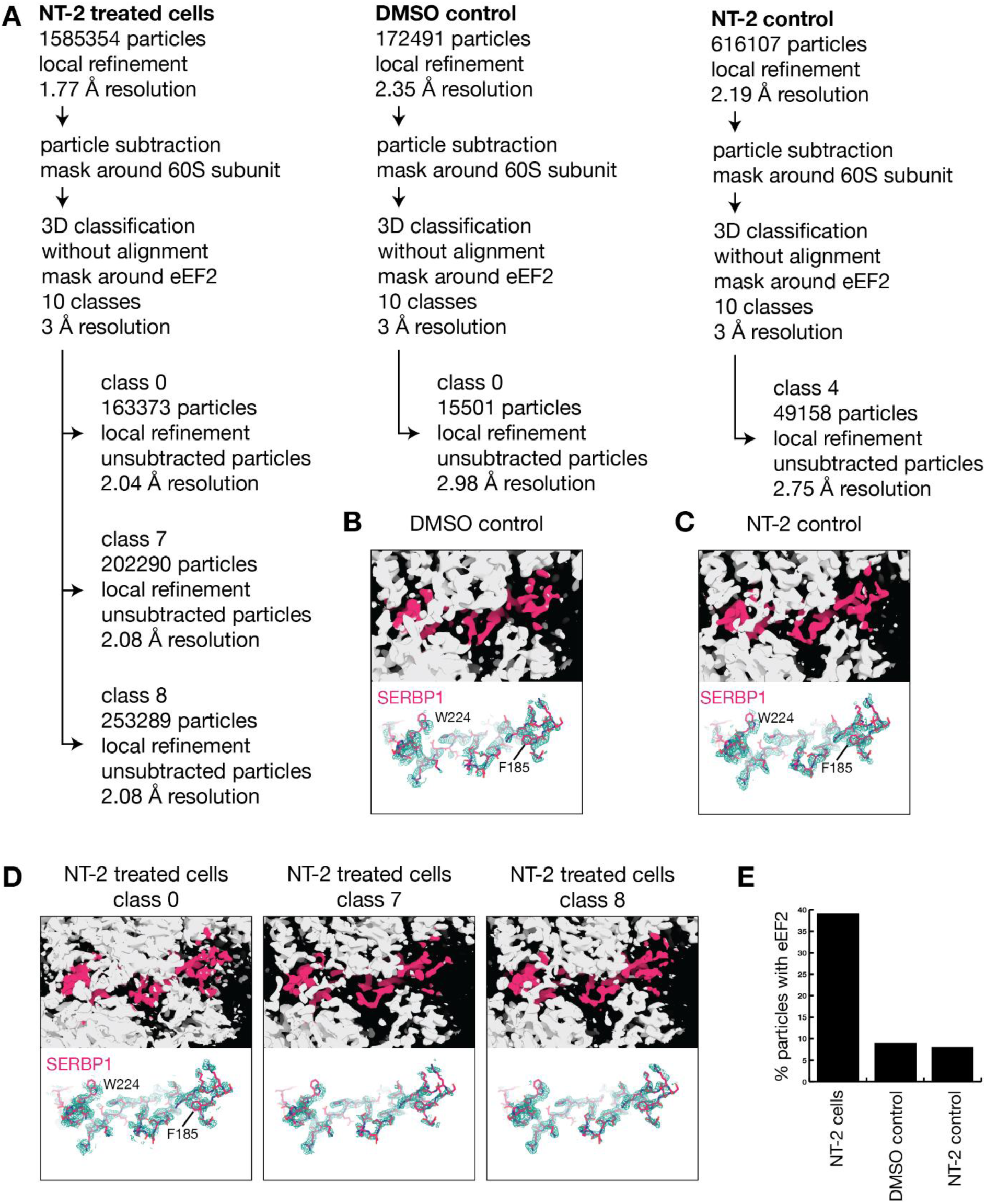
Investigation of eEF2-bound 80S complexes by focused classification. **(A)** Schematic representation of the processing workflow. **(B)** Observed Cryo-EM density for the DMSO control. Ribosome density is shown as white surface and density corresponding to SERBP1 is shown in magenta (top). A rigid-body fitted atomic model of SERBP1 is overlayed with the experimental density of SERBP1 in the DMSO control (bottom). **(C)** Observed Cryo-EM density for the NT-2 control **(D)** Observed Cryo-EM density for the NT-2 treated cells. Density from three classes is shown; all classes contain bound SERBP1. **(E)** Comparison of class sizes after focused classification. While 39% of ribosomes from NT-2 treated cells are found in the dormant SERBP1 and eEF2-bound 80S state, only 9% and 8%, respectively of control ribosomes adopt this state.

**Sup. Table 1.**
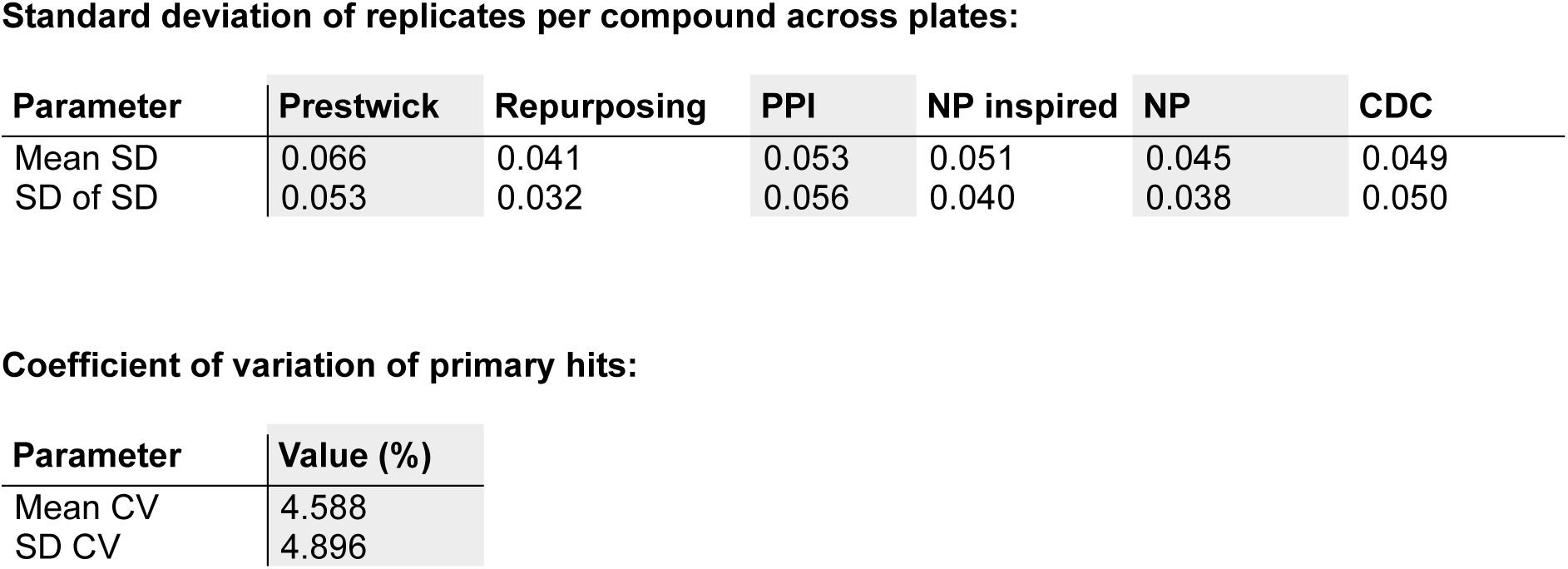
Primary screening reproducibility:

**Sup. Table 2.**
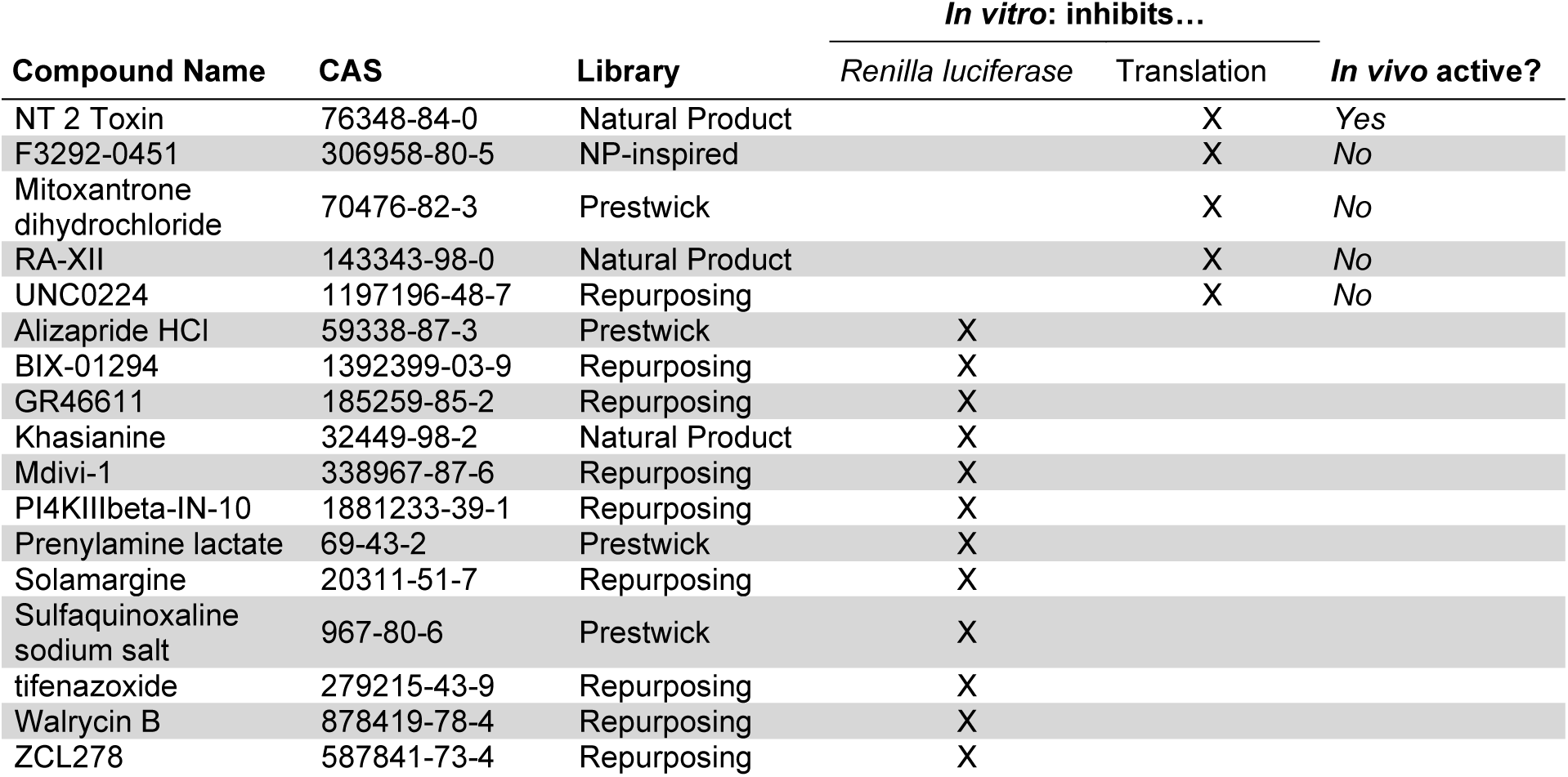
DRC’s secondary hits:

**Sup. Table 3.**
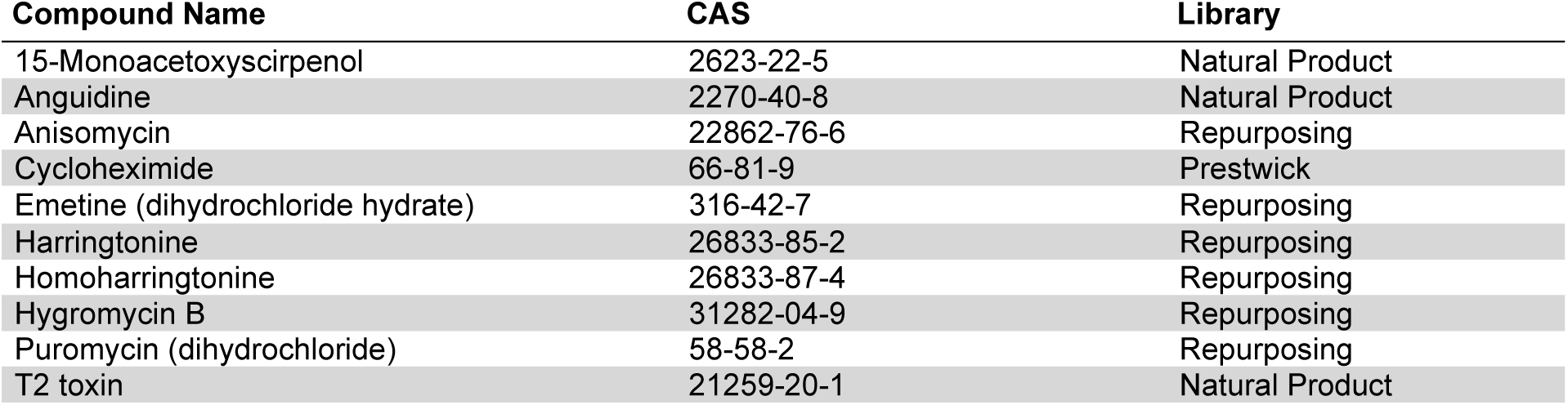
Known translation inhibitors in RC’s secondary hits:

**Sup. Table 4.**
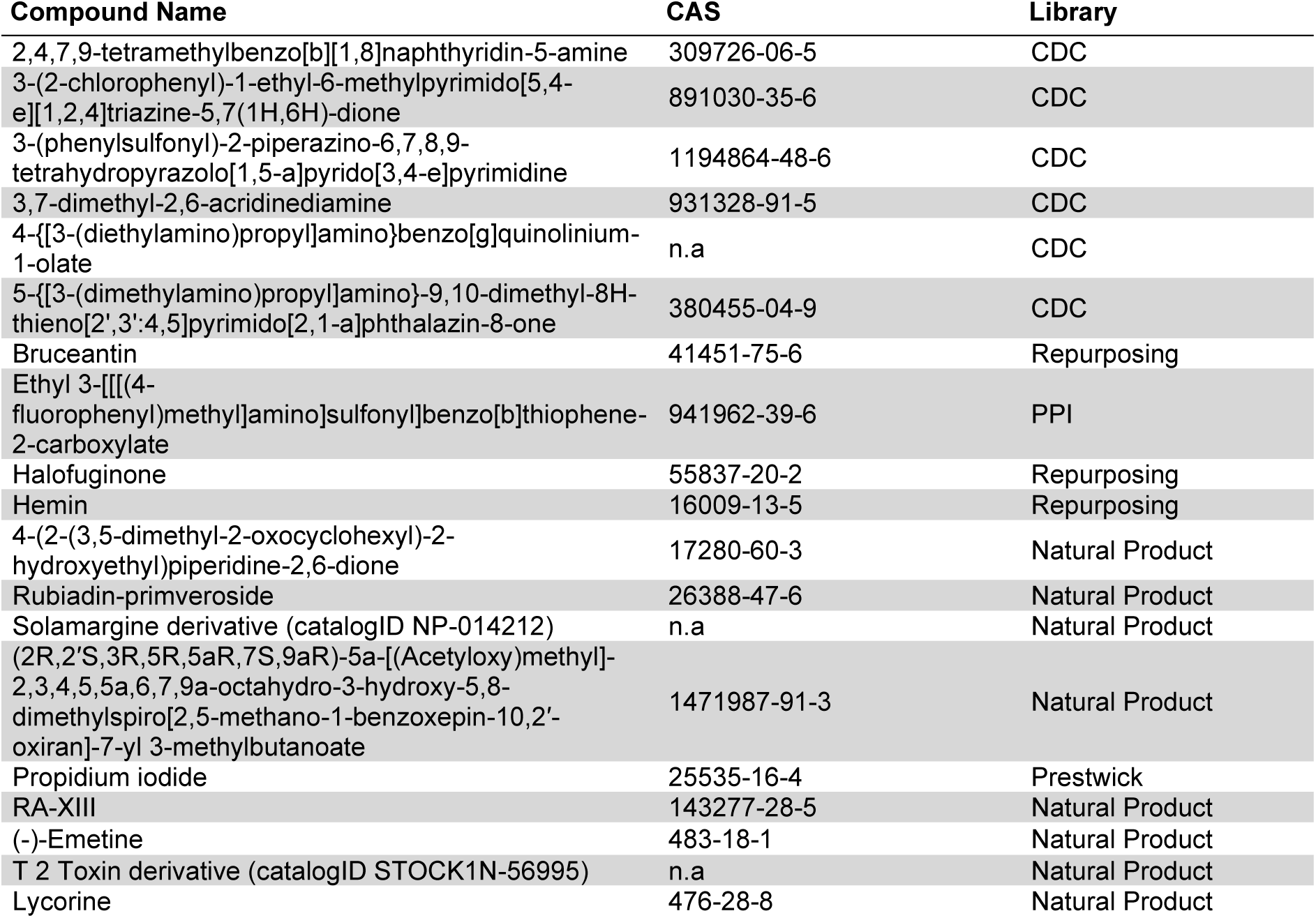
DRC’s secondary hits without further characterization:

**Sup. Table 5.**
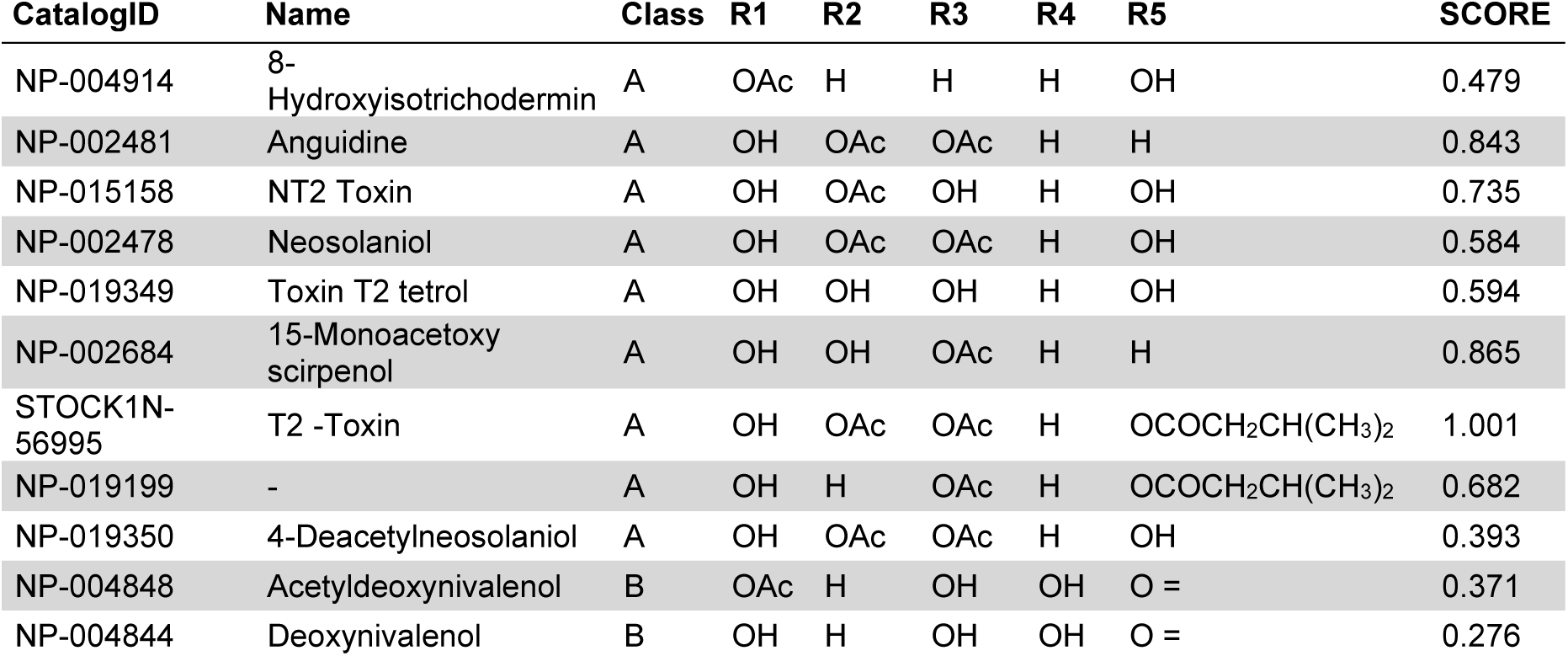
Structural comparison of trichothecenes in screen:

**Sup. Table 6.**
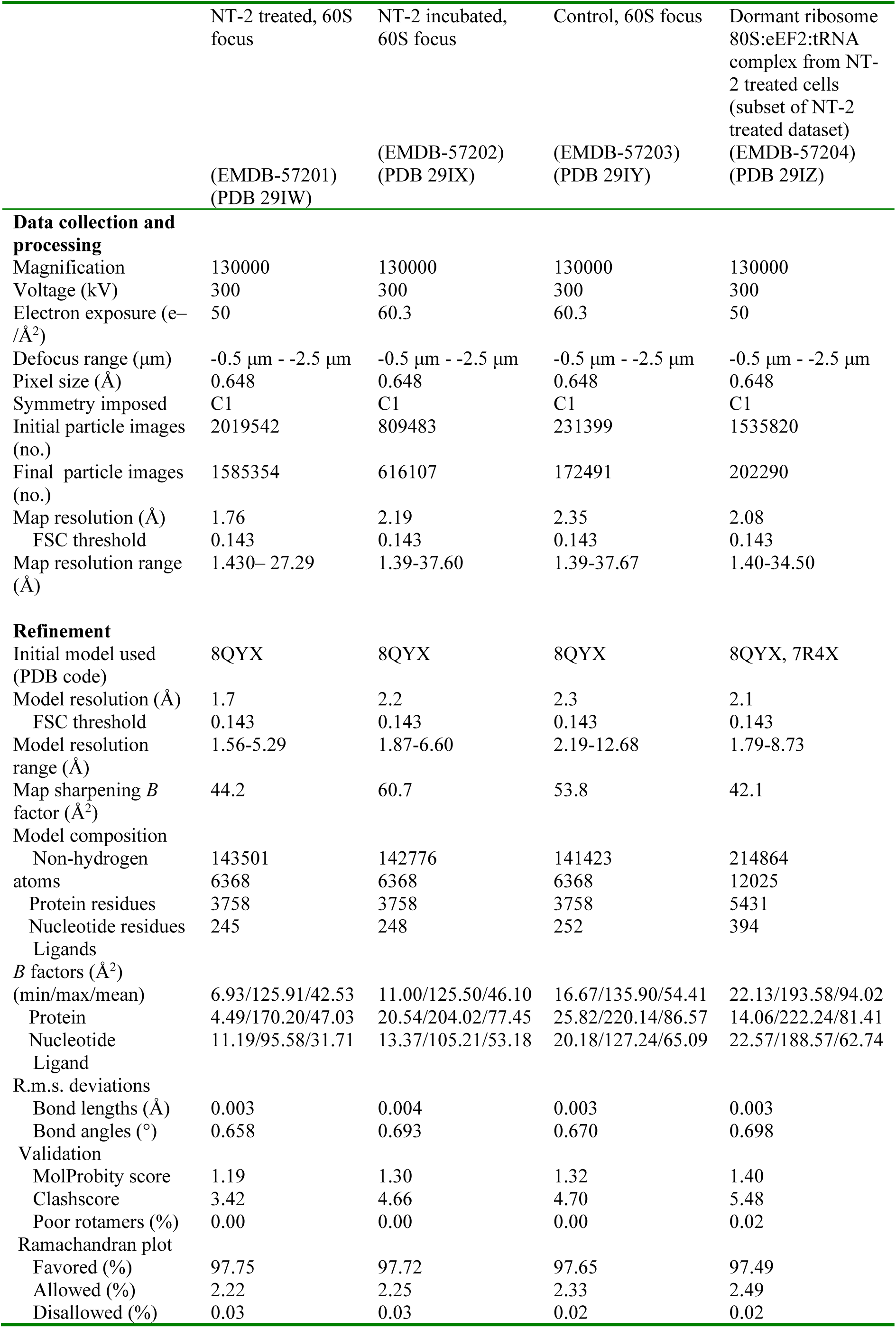
Cryo-EM data collection, refinement and validation statistics.

|  | NT-2 treated, 60S focus | NT-2 incubated, 60S focus | Control, 60S focus | Dormant ribosome 80S:eEF2:tRNA complex from NT-2 treated cells (subset of NT-2 treated dataset) |
| --- | --- | --- | --- | --- |
|  | (EMDB-57201) (PDB 29IW) | (EMDB-57202) (PDB 29IX) | (EMDB-57203) (PDB 29IY) | (EMDB-57204) (PDB 29IZ) |
| **Data collection and processing** |  |  |  |  |
| Magnification | 130000 | 130000 | 130000 | 130000 |
| Voltage (kV) | 300 | 300 | 300 | 300 |
| Electron exposure (e<sup>-</sup>/Å<sup>2</sup>) | 50 | 60.3 | 60.3 | 50 |
| Defocus range (μm) | -0.5 μm - -2.5 μm | -0.5 μm - -2.5 μm | -0.5 μm - -2.5 μm | -0.5 μm - -2.5 μm |
| Pixel size (Å) | 0.648 | 0.648 | 0.648 | 0.648 |
| Symmetry imposed | C1 | C1 | C1 | C1 |
| Initial particle images (no.) | 2019542 | 809483 | 231399 | 1535820 |
| Final particle images (no.) | 1585354 | 616107 | 172491 | 202290 |
| Map resolution (Å) | 1.76 | 2.19 | 2.35 | 2.08 |
| FSC threshold | 0.143 | 0.143 | 0.143 | 0.143 |
| Map resolution range (Å) | 1.430–27.29 | 1.39–37.60 | 1.39–37.67 | 1.40–34.50 |
| **Refinement** |  |  |  |  |
| Initial model used (PDB code) | 8QYX | 8QYX | 8QYX | 8QYX, 7R4X |
| Model resolution (Å) | 1.7 | 2.2 | 2.3 | 2.1 |
| FSC threshold | 0.143 | 0.143 | 0.143 | 0.143 |
| Model resolution range (Å) | 1.56–5.29 | 1.87–6.60 | 2.19–12.68 | 1.79–8.73 |
| Map sharpening *B* factor (Å<sup>2</sup>) | 44.2 | 60.7 | 53.8 | 42.1 |
| Model composition |  |  |  |  |
| Non-hydrogen atoms | 143501 | 142776 | 141423 | 214864 |
| Protein residues | 6368 | 6368 | 6368 | 12025 |
| Nucleotide residues | 3758 | 3758 | 3758 | 5431 |
| Ligands | 245 | 248 | 252 | 394 |
| *B* factors (Å<sup>2</sup>) (min/max/mean) |  |  |  |  |
| Protein | 6.93/125.91/42.53 | 11.00/125.50/46.10 | 16.67/135.90/54.41 | 22.13/193.58/94.02 |
| Nucleotide | 4.49/170.20/47.03 | 20.54/204.02/77.45 | 25.82/220.14/86.57 | 14.06/222.24/81.41 |
| Ligand | 11.19/95.58/31.71 | 13.37/105.21/53.18 | 20.18/127.24/65.09 | 22.57/188.57/62.74 |
| R.m.s. deviations |  |  |  |  |
| Bond lengths (Å) | 0.003 | 0.004 | 0.003 | 0.003 |
| Bond angles (°) | 0.658 | 0.693 | 0.670 | 0.698 |
| Validation |  |  |  |  |
| MolProbity score | 1.19 | 1.30 | 1.32 | 1.40 |
| Clashscore | 3.42 | 4.66 | 4.70 | 5.48 |
| Poor rotamers (%) | 0.00 | 0.00 | 0.00 | 0.02 |
| Ramachandran plot |  |  |  |  |
| Favored (%) | 97.75 | 97.72 | 97.65 | 97.49 |
| Allowed (%) | 2.22 | 2.25 | 2.33 | 2.49 |
| Disallowed (%) | 0.03 | 0.03 | 0.02 | 0.02 |

