## Supplemental information for "A human cell-free translation screen identifies the NT-2 mycotoxin as a ribosomal inhibitor that binds the peptidyl transferase center"

### Supplementary Material

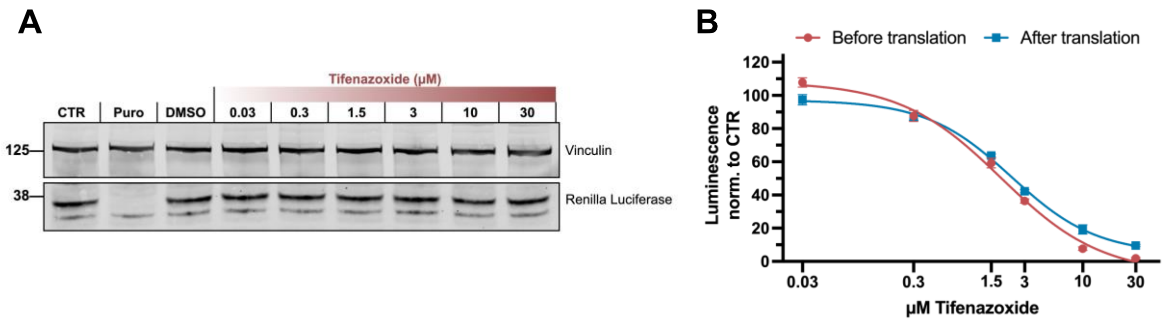


**Supplementary Figure 1 | Characterization of Tifenazoxide as a *Renilla* luciferase inhibitor**
**(A)** Western blot of an *in vitro* translation assay supplemented with tifenazoxide. Cell-free translation reactions were performed with *Renilla* luciferase mRNA in HeLa S3 lysate with increasing Tifenazoxide concentrations (30 nM - 30 µM) and were analyzed by SDS-PAGE and immunoblotting for *Renilla* luciferase (translation product) and Vinculin (loading control). **(B)** Cell-free translation reactions as described in (A) were performed with Tifenazoxide supplementation either before (red) or after (blue) translation. The measured luminescence was normalized to the negative control containing no supplement (CTR). Data are presented as mean values of three biological replicates (sets of translation reactions) averaged after three measurements ± SD.

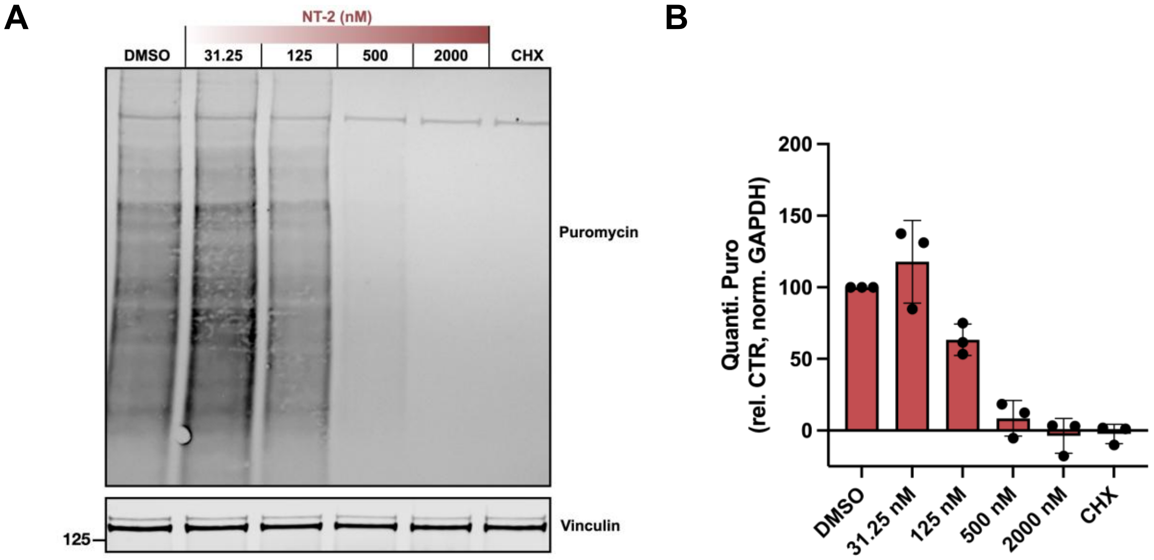


**Supplementary Figure 2 | Effect of NT-2 on translation rates in *BHK-21 cells***
**(A)** Puromycin incorporation assay in BHK-21 cells treated with increasing NT-2 concentrations, DMSO, or CHX. Equal amounts of cell lysate were loaded into a 4-12% Bis-Tris gel and analysed by Western blot probing with anti-puromycin and anti-GAPDH antibodies. **(B)** Quantification of protein synthesis based on signal intensity in the anti-Puromycin-stained blot (Sup Fig 3A) normalized with GAPDH. Bars represent average of three biological replicates (dots) ± SD.

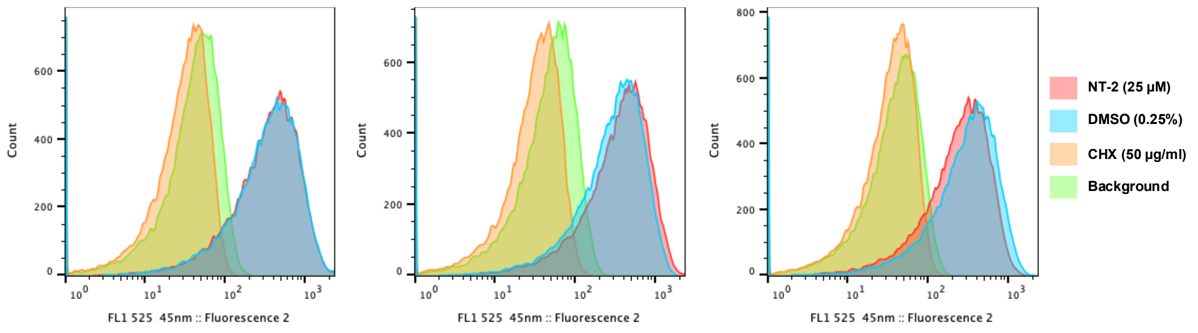


**Supplementary Figure 3 | Effect of NT-2 on translation rates in *S. cerevisiae***
Fluorescence intensity of AHA-labeled yeast cells under different treatments. Cells were treated with NT-2 (25 µM), DMSO (0.25 %, control), CHX (50 µg/mL), or incubated in complete YPD medium (background control) and processed as described in Materials & Methods. Histograms show the distribution of single-cell fluorescence measured by flow cytometry of the three biological replicates.

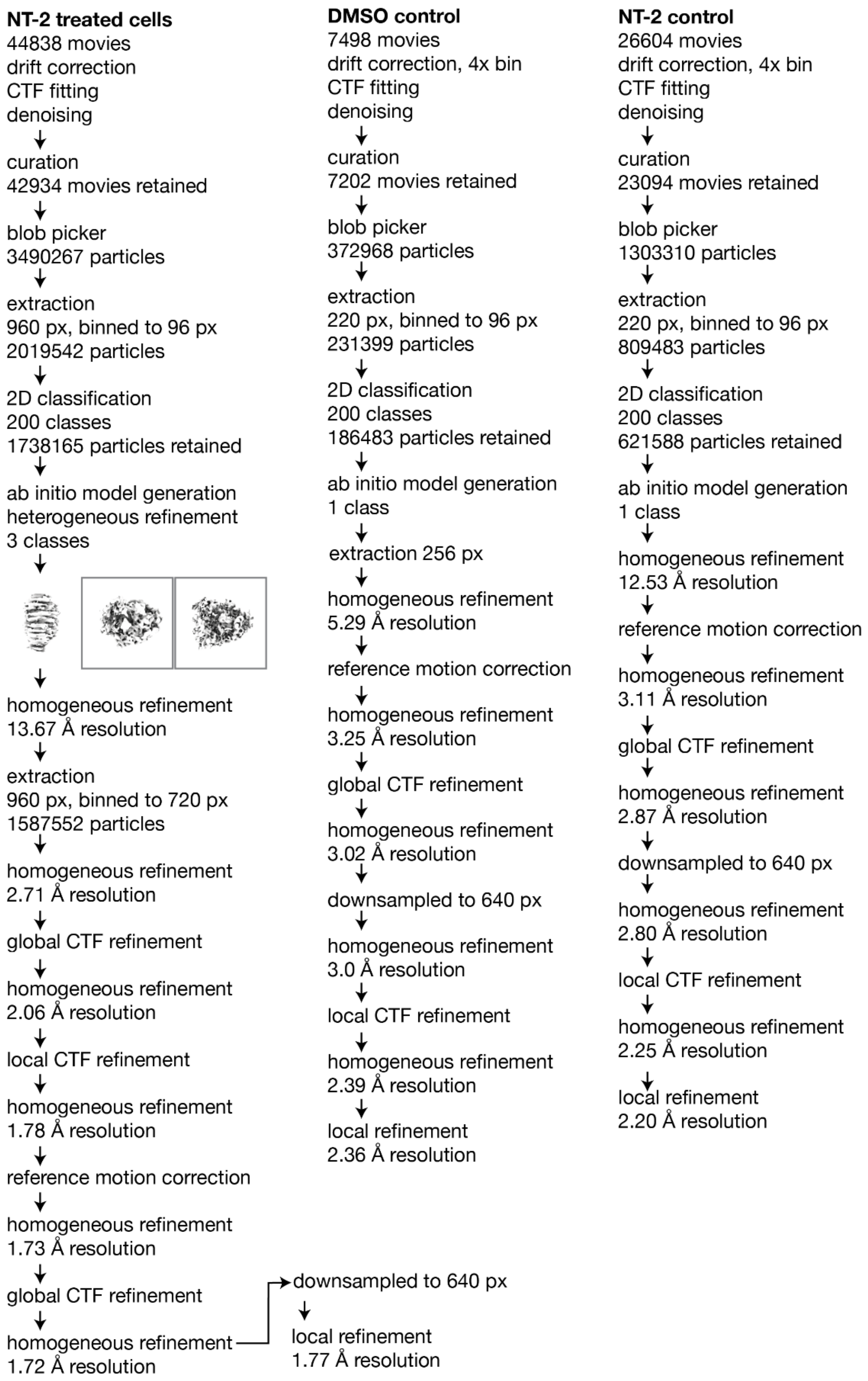


**Supplementary Figure 4 | Cryo-EM data processing flowcharts.**
Image processing workflows for the three datasets: 80S ribosomes isolated from NT-2–treated cells, 80S isolated from DMSO-treated (CTR) cells and NT-2–incubated 80S ribosomes. Flowcharts summarize particle selection, classification, and refinement steps leading to the final reconstructions.

(A)

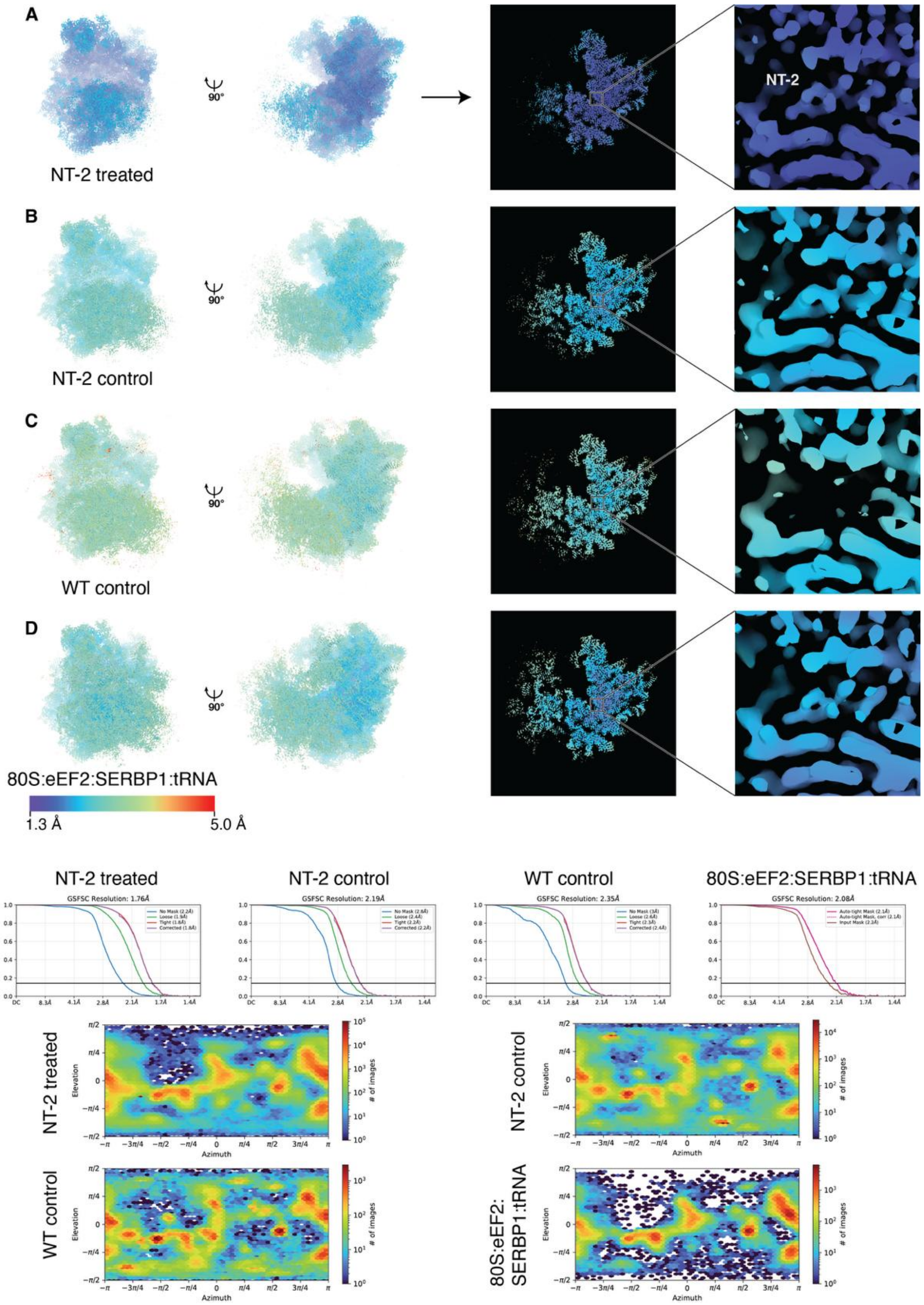


**Supplementary Figure 5 | Resolution maps, FSC curves and particle orientation distribution plots.** Resolution map of monosomes from NT-2 treated cells, focused on the 60S subunit. A resolution map of the surface of the molecule is shown on the left two panels. The center right panel shows a section through the map around the binding site of NT-2. On the right panel, NT-2 and the region directly adjacent are shown. All resolution maps are shown on the same scale (see inset below panel D). **(B)** Resolution map of purified monosomes incubated with NT-2 before vitrification, focused on the 60S subunit. **(C)** Resolution map of purified monosomes, focused on the 60S subunit. NT-2 is absent from the binding site. **(D)** Resolution map of dormant 80S:eEF2:SERBP1:tRNA complex purified from NT-2-treated cells. **(E)** Fourier shell correlation curves. F) Particle distribution plots.

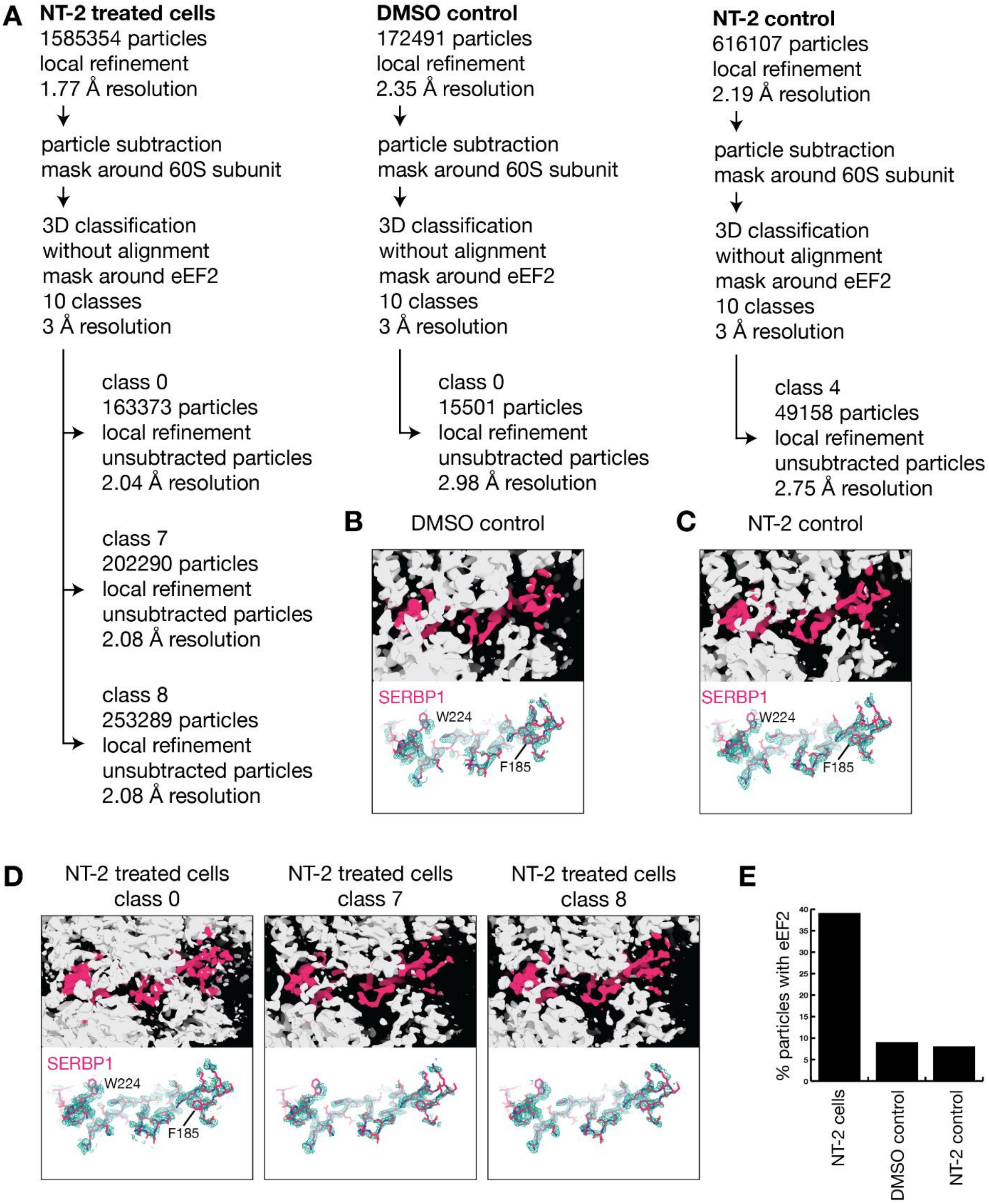


**Supplementary Figure 6| Investigation of eEF2-bound 80S complexes by focused classification.** **(A)** Schematic representation of the processing workflow. **(B)** Observed Cryo-EM density for the DMSO control. Ribosome density is shown as white surface and density corresponding to SERBP1 is shown in magenta (top). A rigid-body fitted atomic model of SERBP1 is overlayed with the experimental density of SERBP1 in the DMSO control (bottom). **(C)** Observed Cryo-EM density for the NT-2 control **(D)** Observed Cryo-EM density for the NT-2 treated cells. Density from three classes is shown; all classes contain bound SERBP1. **(E)** Comparison of class sizes after focused classification. While 39% of ribosomes from NT-2 treated cells are found in the dormant SERBP1 and eEF2-bound 80S state, only 9% and 8%, respectively of control ribosomes adopt this state.

**Sup. Table 1 | Primary screening reproducibility:**

**Standard deviation of replicates per compound across plates:**

| Parameter | Prestwick | Repurposing | PPI | NP inspired | NP | CDC |
|---|---|---|---|---|---|---|
| Mean SD | 0.066 | 0.041 | 0.053 | 0.051 | 0.045 | 0.049 |
| SD of SD | 0.053 | 0.032 | 0.056 | 0.040 | 0.038 | 0.050 |

**Coefficient of variation of primary hits:**

| Parameter | Value (%) |
|---|---|
| Mean CV | 4.588 |
| SD CV | 4.896 |

**Sup. Table 2 | DRC's secondary hits:**

| Compound Name | CAS | Library | *In vitro*: inhibits… | | *In vivo* active? |
|---|---|---|---|---|---|
| | | | *Renilla luciferase* | Translation | |
| NT 2 Toxin | 76348-84-0 | Natural Product | | X | *Yes* |
| F3292-0451 | 306958-80-5 | NP-inspired | | X | *No* |
| Mitoxantrone dihydrochloride | 70476-82-3 | Prestwick | | X | *No* |
| RA-XII | 143343-98-0 | Natural Product | | X | *No* |
| UNC0224 | 1197196-48-7 | Repurposing | | X | *No* |
| Alizapride HCl | 59338-87-3 | Prestwick | X | | |
| BIX-01294 | 1392399-03-9 | Repurposing | X | | |
| GR46611 | 185259-85-2 | Repurposing | X | | |
| Khasianine | 32449-98-2 | Natural Product | X | | |
| Mdivi-1 | 338967-87-6 | Repurposing | X | | |
| PI4KIIIbeta-IN-10 | 1881233-39-1 | Repurposing | X | | |
| Prenylamine lactate | 69-43-2 | Prestwick | X | | |
| Solamargine | 20311-51-7 | Repurposing | X | | |
| Sulfaquinoxaline sodium salt | 967-80-6 | Prestwick | X | | |
| tifenazoxide | 279215-43-9 | Repurposing | X | | |
| Walrycin B | 878419-78-4 | Repurposing | X | | |
| ZCL278 | 587841-73-4 | Repurposing | X | | |

**Sup. Table 3 | Known translation inhibitors in DRC's secondary hits:**

| Compound Name | CAS | Library |
|---|---|---|
| 15-Monoacetoxyscirpenol | 2623-22-5 | Natural Product |
| Anguidine | 2270-40-8 | Natural Product |
| Anisomycin | 22862-76-6 | Repurposing |
| Cycloheximide | 66-81-9 | Prestwick |
| Emetine (dihydrochloride hydrate) | 316-42-7 | Repurposing |
| Harringtonine | 26833-85-2 | Repurposing |
| Homoharringtonine | 26833-87-4 | Repurposing |
| Hygromycin B | 31282-04-9 | Repurposing |
| Puromycin (dihydrochloride) | 58-58-2 | Repurposing |
| T2 toxin | 21259-20-1 | Natural Product |

**Sup. Table 4 | DRC's secondary hits without further characterization:**

| Compound Name | CAS | Library |
|---|---|---|
| 2,4,7,9-tetramethylbenzo[b][1,8]naphthyridin-5-amine | 309726-06-5 | CDC |
| 3-(2-chlorophenyl)-1-ethyl-6-methylpyrimido[5,4-e][1,2,4]triazine-5,7(1H,6H)-dione | 891030-35-6 | CDC |
| 3-(phenylsulfonyl)-2-piperazino-6,7,8,9-tetrahydropyrazolo[1,5-a]pyrido[3,4-e]pyrimidine | 1194864-48-6 | CDC |
| 3,7-dimethyl-2,6-acridinediamine | 931328-91-5 | CDC |
| 4-{[3-(diethylamino)propyl]amino}benzo[g]quinolinium-1-olate | n.a | CDC |
| 5-{[3-(dimethylamino)propyl]amino}-9,10-dimethyl-8H-thieno[2',3':4,5]pyrimido[2,1-a]phthalazin-8-one | 380455-04-9 | CDC |
| Bruceantin | 41451-75-6 | Repurposing |
| Ethyl 3-[[[(4-fluorophenyl)methyl]amino]sulfonyl]benzo[b]thiophene-2-carboxylate | 941962-39-6 | PPI |
| Halofuginone | 55837-20-2 | Repurposing |
| Hemin | 16009-13-5 | Repurposing |
| 4-(2-(3,5-dimethyl-2-oxocyclohexyl)-2-hydroxyethyl)piperidine-2,6-dione | 17280-60-3 | Natural Product |
| Rubiadin-primveroside | 26388-47-6 | Natural Product |
| Solamargine derivative (catalogID NP-014212) | n.a | Natural Product |
| (2R,2′S,3R,5R,5aR,7S,9aR)-5a-[(Acetyloxy)methyl]-2,3,4,5,5a,6,7,9a-octahydro-3-hydroxy-5,8-dimethylspiro[2,5-methano-1-benzoxepin-10,2′-oxiran]-7-yl 3-methylbutanoate | 1471987-91-3 | Natural Product |
| Propidium iodide | 25535-16-4 | Prestwick |
| RA-XIII | 143277-28-5 | Natural Product |
| (-)-Emetine | 483-18-1 | Natural Product |
| T 2 Toxin derivative (catalogID STOCK1N-56995) | n.a | Natural Product |
| Lycorine | 476-28-8 | Natural Product |

**Sup. Table 5 | Structural comparison of trichothecenes in screen:**

| CatalogID | Name | Class | R1 | R2 | R3 | R4 | R5 | SCORE |
|---|---|---|---|---|---|---|---|---|
| NP-004914 | 8-Hydroxyisotrichodermin | A | OAc | H | H | H | OH | 0.479 |
| NP-002481 | Anguidine | A | OH | OAc | OAc | H | H | 0.843 |
| NP-015158 | NT2 Toxin | A | OH | OAc | OH | H | OH | 0.735 |
| NP-002478 | Neosolaniol | A | OH | OAc | OAc | H | OH | 0.584 |
| NP-019349 | Toxin T2 tetrol | A | OH | OH | OH | H | OH | 0.594 |
| NP-002684 | 15-Monoacetoxy scirpenol | A | OH | OH | OAc | H | H | 0.865 |
| STOCK1N-56995 | T2 -Toxin | A | OH | OAc | OAc | H | $OCOCH_2CH(CH_3)_2$ | 1.001 |
| NP-019199 | - | A | OH | H | OAc | H | $OCOCH_2CH(CH_3)_2$ | 0.682 |
| NP-019350 | 4-Deacetylneosolaniol | A | OH | OAc | OAc | H | OH | 0.393 |
| NP-004848 | Acetyldeoxynivalenol | B | OAc | H | OH | OH | O = | 0.371 |
| NP-004844 | Deoxynivalenol | B | OH | H | OH | OH | O = | 0.276 |

| **Sup. Table 6 \| Cryo-EM data collection, refinement and validation statistics** | NT-2 treated, 60S focus (EMDB-57201) (PDB 29IW) | NT-2 incubated, 60S focus (EMDB-57202) (PDB 29IX) | Control, 60S focus (EMDB-57203) (PDB 29IY) | Dormant ribosome 80S:eEF2:tRNA complex from NT-2 treated cells (subset of NT-2 treated dataset) (EMDB-57204) (PDB 29IZ) |
|---|---|---|---|---|
| **Data collection and processing** | | | | |
| Magnification | 130000 | 130000 | 130000 | 130000 |
| Voltage (kV) | 300 | 300 | 300 | 300 |
| Electron exposure (e–/Å$^2$) | 50 | 60.3 | 60.3 | 50 |
| Defocus range (µm) | -0.5 µm - -2.5 µm | -0.5 µm - -2.5 µm | -0.5 µm - -2.5 µm | -0.5 µm - -2.5 µm |
| Pixel size (Å) | 0.648 | 0.648 | 0.648 | 0.648 |
| Symmetry imposed | C1 | C1 | C1 | C1 |
| Initial particle images (no.) | 2019542 | 809483 | 231399 | 1535820 |
| Final particle images (no.) | 1585354 | 616107 | 172491 | 202290 |
| Map resolution (Å) | 1.76 | 2.19 | 2.35 | 2.08 |
| FSC threshold | 0.143 | 0.143 | 0.143 | 0.143 |
| Map resolution range (Å) | 1.430– 27.29 | 1.39-37.60 | 1.39-37.67 | 1.40-34.50 |
| **Refinement** | | | | |
| Initial model used (PDB code) | 8QYX | 8QYX | 8QYX | 8QYX, 7R4X |
| Model resolution (Å) | 1.8 | 2.2 | 2.3 | 2.1 |
| FSC threshold | 0.143 | 0.143 | 0.143 | 0.143 |
| Model resolution range (Å) | 1.56-5.29 | 1.87-6.60 | 2.19-12.68 | 1.79-8.73 |
| Map sharpening *B* factor (Å$^2$) | 44.2 | 60.7 | 53.8 | 42.1 |
| Model composition | | | | |
| Non-hydrogen atoms | 143501 | 142776 | 141423 | 214864 |
| Protein residues | 6368 | 6368 | 6368 | 12025 |
| Nucleotide residues | 3758 | 3758 | 3758 | 5431 |
| Ligands | 245 | 248 | 252 | 394 |
| *B* factors (Å$^2$) (min/max/mean) | | | | |
| Protein | 0.00/107.06/22.11 | 11.00/125.50/46.10 | 16.67/135.90/54.41 | 0.53/137.45/41.49 |
| Nucleotide | 1.74/166.41/43.37 | 20.54/204.02/77.45 | 25.82/220.14/86.57 | 0.77/220.13/72.37 |
| Ligand | 3.54/98.13/26.98 | 13.37/105.21/53.18 | 20.18/127.24/65.09 | 0.00/166.36/55.96 |
| R.m.s. deviations | | | | |
| Bond lengths (Å) | 0.003 | 0.004 | 0.003 | 0.003 |
| Bond angles (°) | 0.679 | 0.693 | 0.670 | 0.685 |
| Validation | | | | |
| MolProbity score | 1.23 | 1.30 | 1.32 | 1.43 |
| Clashscore | 3.78 | 4.66 | 4.70 | 5.79 |
| Poor rotamers (%) | 0.00 | 0.00 | 0.00 | 0.02 |
| Ramachandran plot | | | | |
| Favored (%) | 97.72 | 97.72 | 97.65 | 97.40 |
| Allowed (%) | 2.23 | 2.25 | 2.33 | 2.60 |
| Disallowed (%) | 0.05 | 0.03 | 0.02 | 0.01 |
